# Human monoclonal antibodies against *Staphylococcus aureus* surface antigens recognize *in vitro* biofilm and *in vivo* implant infections

**DOI:** 10.1101/2021.02.09.429966

**Authors:** Lisanne de Vor, Bruce van Dijk, Kok P.M. van Kessel, Jeffrey S. Kavanaugh, Carla J.C. de Haas, Piet C. Aerts, Marco C. Viveen, Edwin C.H. Boel, Ad C. Fluit, Jakub M. Kwiecinski, Gerard C. Krijger, Ruud M. Ramakers, Freek J. Beekman, Ekaterina Dadachova, Marnix G.E.H. Lam, H. Charles Vogely, Bart C.H. van der Wal, Jos A.G. van Strijp, Alexander R. Horswill, Harrie Weinans, Suzan H.M. Rooijakkers

## Abstract

Implant-associated *Staphylococcus aureus* infections are difficult to treat because of biofilm formation. Bacteria in a biofilm are often insensitive to antibiotics and host immunity. Monoclonal antibodies (mAbs) could provide an alternative approach to improve the diagnosis and/or treatment of biofilm-related infections. Here we show that mAbs targeting common surface components of *S. aureus* can recognize clinically relevant biofilm types. We identify two groups of antibodies: one group that uniquely binds *S. aureus* in biofilm state and one that recognizes *S. aureus* in both biofilm and planktonic state. In a mouse model, we show that mAb 4497 (recognizing wall teichoic acid (WTA)) specifically localizes to biofilm-infected implants. In conclusion, we demonstrate the capacity of several human mAbs to detect *S. aureus* biofilms *in vitro* and *in vivo*. This is an important first step to develop mAbs for imaging or treating *S. aureus* biofilms.

## Introduction

Implant-related infections are difficult to treat because of the ability of many bacterial species to form biofilm (1). Biofilms are bacterial communities that adhere to abiotic surfaces (such as medical implants) using a self-made extracellular polymeric substance (EPS), consisting of proteins, polysaccharides and extracellular DNA(2, 3). Bacteria in a biofilm are physically different from planktonic (free floating) bacteria and often more tolerant to antibiotics (4). For instance, the EPS forms an important penetration barrier for many antimicrobial agents (2, 5). In addition, most antibiotics cannot kill bacteria in a biofilm, because they are in a metabolically inactive state (6) and thus resistant to the antibiotics that act on active cellular processes (such as transcription/translation or cell wall formation(7)). Another complication is that biofilm infections often occur in areas of the body that are not easily accessible for treatment without invasive surgical procedures. Consequently, treatment consists of long-term antibiotic regimens or replacement of the infected implant. Specific and noninvasive laboratory tests for early detection are not yet available and the diagnosis is often made only at advanced stages. This failure to detect biofilms adds further complications to effective diagnosis and treatment of these infections.

The human pathogen *S. aureus* is the leading cause of healthcare associated infections (8, 9). Today, 25% of healthcare-associated infections are related to medical implants such as heart valves, intravenous catheters and prosthetic joints (10). *S. aureus* causes one third of all implant-related infections in Europe and the United States (11, 12) and is known for its ability to form biofilm (1). Due to the absence of a vaccine and the emergence of methicillin-resistant *S. aureus* (MRSA), there is a clear need for diagnostic tools and alternative therapies for *S. aureus* biofilm infections.

Antibody-based biologicals could provide an alternative approach to improve the diagnosis and/or treatment of *S. aureus* biofilm-related infections. Monoclonal antibodies (mAb) may be exploited as vehicles to specifically bring anti-biofilm agents (such as radionuclides, enzymes or photosensitizers) to the site of infection (13–20). Furthermore, radioactively labeled mAbs could be used for early diagnosis of biofilm related infections. At present, only one mAb recognizing *S. aureus* biofilm has been identified. This F598 antibody recognizes poly-*N*-acetyl glucosamine (PNAG) (21, 22) (also known as polysaccharide intercellular adhesion (PIA) (23, 24)), a highly positively charged polysaccharide that was first recognized as a major EPS component of *S. aureus* biofilm. However, PNAG is not the only component of *S. aureus* biofilms. Recently, it has become clear that *S. aureus* may also use cell wall anchored proteins and eDNA to facilitate initial attachment and intercellular adhesion (25–28). In fact, deletion of the *icaADBC* locus (encoding PNAG) does not impair biofilm formation in multiple *S. aureus* strains (4, 28, 29). These biofilms, referred to as PNAG-negative, are phenotypically different from PNAG-positive biofilm (30–35). In this study we show that previously identified mAbs against staphylococcal surface structures are capable of recognizing both PNAG-negative and PNAG-positive *S. aureus* biofilms. Importantly, we show that some of these mAbs recognize *S. aureus* in both biofilm and planktonic state, which is crucial because release and dissemination of planktonic cells from biofilm-infected implants leads to life-threatening complications (36). Finally, using SPECT/CT imaging, we show that radiolabeled mAbs have the potential to visualize implant-associated infections *in vivo.*

## Results

### Production of monoclonal antibodies and validation of S. aureus biofilms

In order to study the reactivity of monoclonal antibodies with *S. aureus* biofilms, we selected mAbs that were previously found to recognize surface components of planktonic *S. aureus* cells(14, 37). Specifically, we generated two antibodies recognizing cell wall teichoic acids (WTA) (4461-IgG and 4497-IgG)(38, 39), one antibody against surface proteins of the SDR family (rF1-IgG)(40), one antibody against Clumping factor A (ClfA)) (T1-2-IgG) and one antibody of which the exact target is yet unknown (CR5132-IgG) **(Fig. 1a)**. As a positive control, we generated F598-IgG against PNAG (21). As negative controls, we produced one antibody recognizing the hapten dinitrophenol (DNP) (G2a-2-IgG)(41) and one recognizing HIV protein gp120 (b12-IgG)(42). The variable heavy and light chain sequences of these antibodies were obtained from different scientific and patent publications **(Table S1)** and cloned into expression vectors to produce full-length human IgG1 (kappa) antibodies in EXPI293F cells.

**Figure 1.**
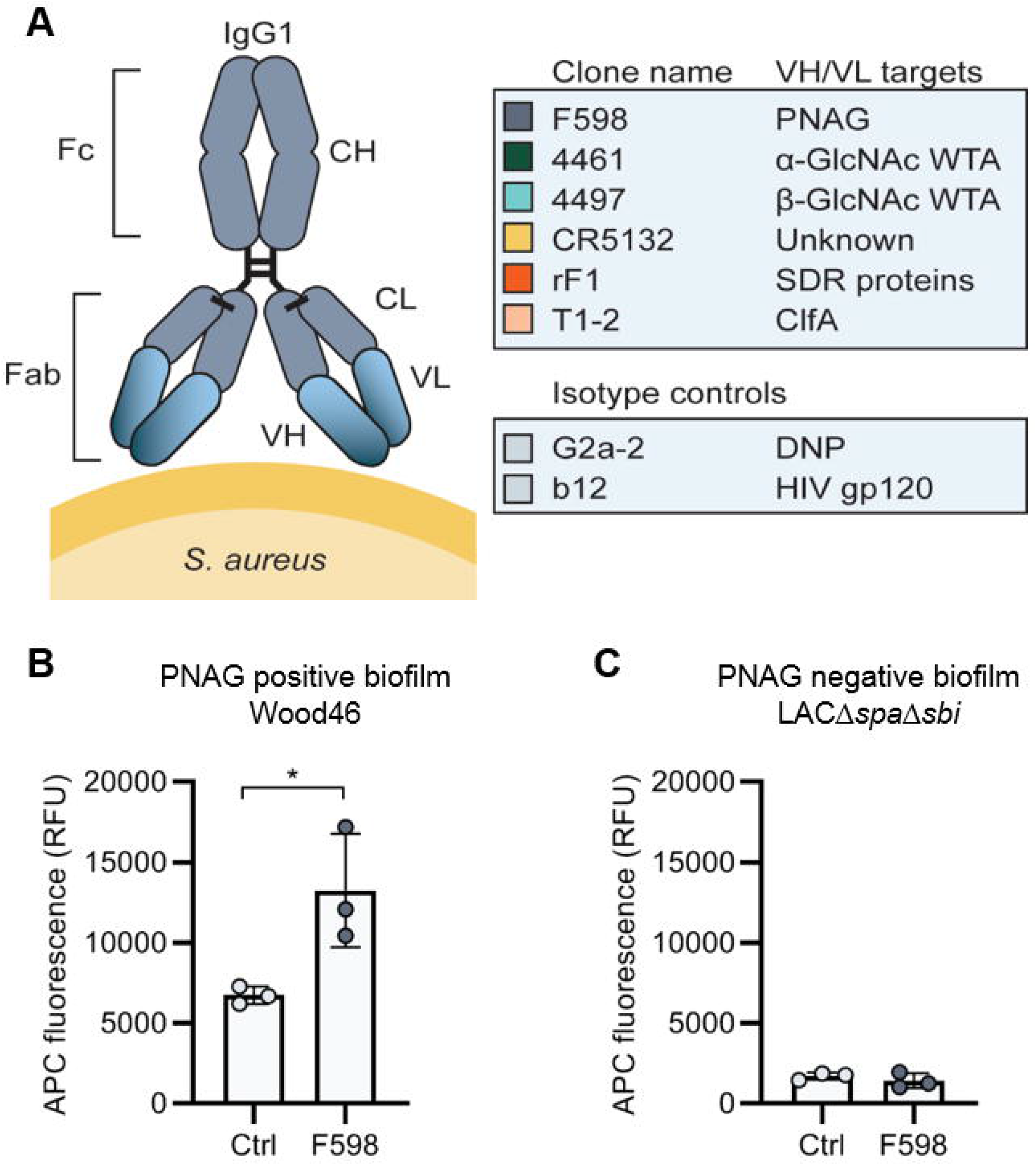
Production of mAbs and validation of biofilm. (A) Human IgG1 antibodies are large (150 kDa) proteins, consisting of two functional domains. The fragment antigen binding (Fab) region confers antigen specificity, while the crystallizable fragment (Fc) region drives interactions with the immune system. Each IgG1 is composed of 2 identical heavy chains and two identical light chains, which all consist of a constant (CH, CL) and a variable (VH, VL) domain. A panel of 6 human IgG1 mAbs that recognize polysaccharide and protein components on the cell surface of *S. aureus* and two non-specific isotype controls was produced. Variable heavy (VH) and light (VL) chain sequences obtained from different scientific and patent publications were cloned in homemade expression vectors containing human heavy chain (HC) and light chain (LC) constant regions, respectively. (B,C) Biofilm of Wood46 (B) and LACΔspaΔsbi (C) were grown for 24h and incubated with 66 nM F598-IgG1. MAb binding was detected using APC-labeled anti-human IgG antibodies and a plate reader and plotted as fluorescence intensity per well. Data represent mean + SD of at least 3 independent experiments. A ratio paired t-test was performed to test for differences in antibody binding versus control and displayed only when significant as *P ≤ 0.05, **P ≤ 0.01, ***P ≤ 0.001, or ****P ≤ 0.0001.

Since we were interested in the reactivity of these mAbs with both PNAG-positive and PNAG-negative biofilms, we selected two *S. aureus* to serve as models for these different biofilm phenotypes. Because PNAG-positive biofilm is able to attach to uncoated polystyrene microtiter plates via electrostatic interactions(43, 44), we screened *S. aureus* strains available in our laboratory for biofilm formation on uncoated plates. This screen revealed strain Wood46 (ATCC 10832) as a robust biofilm former on uncoated, polystyrene microtiter plates **(Fig. S1a)**. This is consistent with the fact that Wood46 is known to produce PNAG (45). An extra advantage of Wood46 is its low surface expression of IgG binding Staphylococcal protein A (SpA), which complicates the detection of antibodies(46, 47). As a model strain for PNAG-negative biofilms we used LAC (48–50), a member of the USA300 lineage which has emerged as the common cause of healthcare-associated MRSA infections, including implant infections (51–54). Previous studies have demonstrated that LAC is capable of forming robust biofilm with no detectable PNAG (15, 28, 30, 55, 56). Here, we used LACΔ*spa*Δ*sbi*, a mutant that lacks both SpA and a second immunoglobulin-binding protein (Sbi). To confirm the EPS composition of Wood46 and LACΔ*spa*Δ*sbi* biofilm, we treated biofilms with different EPS-degrading enzymes that degrade either PNAG (Dispersin B (DspB)) or extracellular DNA (DNaseI). As expected, LACΔ*spa*Δ*sbi* biofilm **(Fig. S1b)** was sensitive to DNase I but not DspB while Wood46 biofilm **(Fig. S1c)** was sensitive to DspB but not DNase I. At ultrastructural level, scanning electron microscopy (SEM) also verified the formation of phenotypically different biofilm by both strains **(Fig. S1de)**. Additionally, we verified that F598-IgG1, the only mAb in our panel that has been reported to bind biofilm(21), indeed recognizes PNAG-positive biofilm of Wood46 **(Fig. 1b, S2)** but not PNAG-negative biofilm of LACΔ*spa*Δ*sbi* **(Fig. 1c, S2)**.

### 4461-IgG1 and 4497-IgG1 against WTA recognize PNAG-positive and PNAG-negative S. aureus biofilm

Next, we tested the binding of other monoclonal antibodies to *S. aureus* biofilms, starting with two well-defined antibodies recognizing WTA, the most abundant glycopolymer on the surface of *S. aureus* (57). MAbs 4461 and 4497 recognize different forms of WTA: while 4461 binds WTA with α-linked GlcNAc, 4497 recognizes β-linked GlcNAc(38, 39).The extent to which WTA is modified with GlcNAc depends both on the presence of enzymes responsible for α- or β-glycosylation (58) and growth conditions. First, we studied binding of 4461-IgG1 and 4497-IgG1 to planktonic cultures of Wood46 and LACΔ*spa*Δ*sbi* **(Fig. 2a)**. In line with the fact that Wood46 is negative for the enzyme responsible for α-GlcNAc glycosylation of WTA (TarM (58)), we observed no binding of 4461-IgG1 to planktonic Wood46. In contrast, 4461-IgG1 bound strongly to planktonic LACΔ*spa*Δ*sbi*. For 4497-IgG, we observed that 4497-IgG1 bound strongly to planktonic Wood46 cells but very weakly to planktonic LACΔ*spa*Δ*sbi* (**Fig. 2a**).

**Figure 2.**
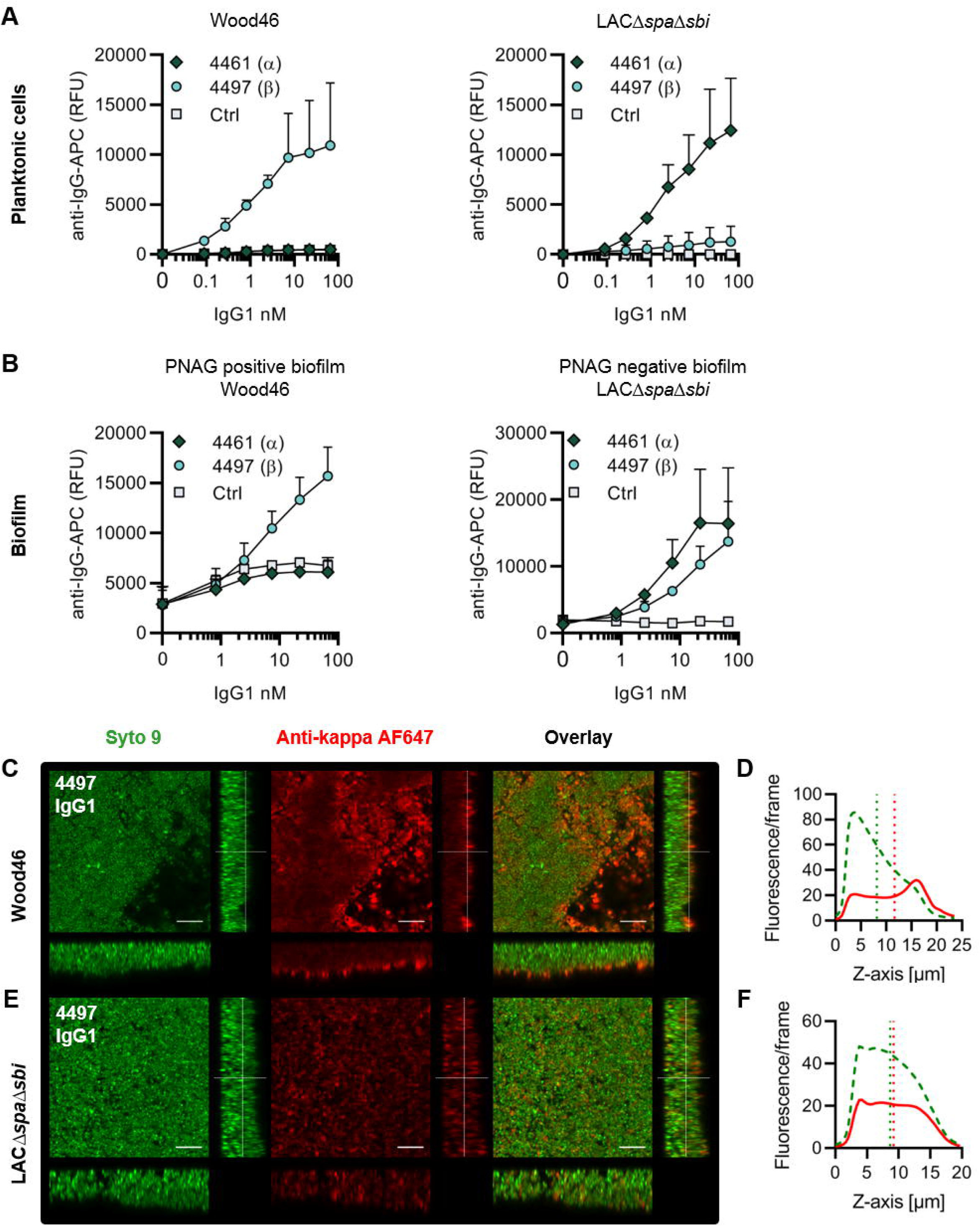
IgG1 mAbs against WTA bind *S. aureus* in planktonic and biofilm mode. (A) Planktonic bacteria of Wood46 (left) and LACΔ*spa*Δ*sbi* (right) were grown to exponential phase and incubated with a concentration range of 4461-IgG1 or 4497-IgG1. MAb binding was detected using APC-labeled anti-human IgG antibodies and flow cytometry and plotted as geoMFI of the bacterial population. (B) Biofilm of Wood46 (left) and LACΔ*spa*Δ*sbi* (right) were grown for 24 h and incubated with a concentration range of 4461-IgG1 or 4497-IgG1. MAb binding was detected using APC-labeled anti-human IgG antibodies and a plate reader and plotted as fluorescence intensity per well. Data represent mean + SD of at least 3 independent experiments. (C,D,E,F) Biofilm was grown for 24 h and incubated with 66 nM IgG1 mAb. Bacteria were visualized by Syto9 (green) and mAb binding was detected by staining with Alexa Fluor 647 conjugated goat-anti-human-kappa F(ab’)2 antibody (red). (C,E) Orthogonal views are representative for a total of three Z-stacks per condition and at least 2 independent experiments. Scale bars: 10 μm. (D,F) Z-stack profile plotting the total fluorescence of Syto9 (green, dotted line) and AF647 (red line) per frame versus the depth (μm) of the corresponding Z-stack. Vertical green and red lines represent the center of mass of the total fluorescent signal.

Upon studying binding of WTA-specific antibodies to biofilms, we observed that 4497-IgG1 strongly bound to PNAG-positive biofilm formed by Wood46 (**Fig. 2b**). While F598-IgG1 exclusively binds PNAG-positive biofilms but not planktonic *S. aureus* **(Fig. S2)**, 4497-IgG1 can bind *S. aureus* in both planktonic and biofilm states. This is important because in the biofilm life cycle, planktonic cells can be released from a biofilm and disseminate to other locations in the body (36). Apart from recognizing PNAG-positive biofilms, 4497-IgG1 also bound the PNAG-negative biofilm formed by LACΔ*spa*Δ*sbi* **(Fig. 2b)**. This is remarkable because 4497-IgG1 did not potently bind planktonic LACΔ*spa*Δ*sbi* **(Fig. 2a)** and suggests that β-glycosylation of WTA is upregulated during LACΔ*spa*Δ*sbi* biofilm formation. Finally, we observe that also 4461-IgG1 effectively recognizes PNAG-negative biofilms. In all, these data identify mAbs against WTA as potent binders of PNAG-positive (4497) and PNAG-negative (4461 and 4497) biofilms.

IgG antibodies are large proteins (150 kD) and therefore penetration into extracellular matrices, like biofilm EPS, can be an issue (59) (60). Besides a penetration barrier, biofilm EPS potentially shields antigens from antibody binding. Using confocal microscopy, we confirmed binding of anti-WTA mAbs and looked at the 3D distribution of IgG1 mAbs in *in vitro* biofilm. Biofilm was cultured in chambered microscopy slides and incubated with IgG1 mAbs or isotype controls **(Fig. S3, S4)**. Bound mAbs were detected by using AF647-labeled anti-human-kappa-antibodies; bacteria were visualized using DNA dye Syto 9. A total of 3 Z-stacks were acquired at random locations in each chamber of the slide. Z-stacks were visualized as orthogonal views + Z-stack profiles, which show the Syto9 and AF647 fluorescence per focal plane. Looking at Wood46 PNAG-positive biofilm **(Fig. 2cd)**, we observed that 4497-IgG1 binding concentrated at the outer biofilm surface. In contrast, we observed that 4497-IgG1, distributed equally across the PNAG-negative biofilm 3D structure **(Fig. 2ef)**, while 4461-IgG1 **(Fig. S3)** shows a trend to binding at the outer surface of PNAG-negative biofilm. Importantly, isotype controls showed no binding **(Fig. S3, S4)**. In conclusion, we show that mAbs recognizing polysaccharides WTA α-GlcNAc and WTA β-GlcNAc are able to bind their targets when bacteria are growing in biofilm mode.

### CR5132-IgG1 discriminates between planktonic bacteria and biofilm

MAb CR5132 was discovered through phage display libraries from human memory B cells (US 2012/0141493 A1) and was selected for binding to staphylococcal colonies scraped from plates. Since such colonies more closely resemble a surface attached biofilm than free floating cells(61), we were curious whether this mAb could recognize biofilm. Intriguingly, CR5132-IgG1 showed almost no detectable binding to planktonic LACΔ*spa*Δ*sbi* or Wood46 (**Fig. 3a)** but it bound strongly to both PNAG-negative and PNAG-positive biofilm formed by these strains **(Fig. 3b)**. This makes CR5132 a unique and interesting mAb since it targets both types of *S. aureus* biofilms and is able to discriminate between planktonic bacteria and biofilm. Confocal microscopy suggests that CR5132-IgG1 penetrates best into the biofilm, as binding of CR5132-IgG1 was distributed more equally across Z-stacks of PNAG-positive biofilm **(Fig. 3cd)** than 4497-IgG1 **(Fig. 2cd)** and other mAbs **(Fig. S4)**. We observed that CR5132-IgG1 also distributed equally across the PNAG-negative biofilm **(Fig. 3ef)**. Because of the interesting binding phenotype of CR5132-IgG1, we performed experiments to identify its target. LTA was originally identified as one of the targets of CR5132 (US 2012/0141493 A1), but the quality of commercial LTA preparations varies greatly and often contains other components (62, 63). Therefore we tested binding to *S. aureus* purified cell wall components LTA and peptidoglycan and we used synthetic α-GlcNAc and β-GlcNAc WTA beads(64) to test binding to pure WTA structures. This way, we identified WTA β-GlcNAc instead of LTA as one of the targets of CR5132 **(Fig. S5)**.

**Figure 3.**
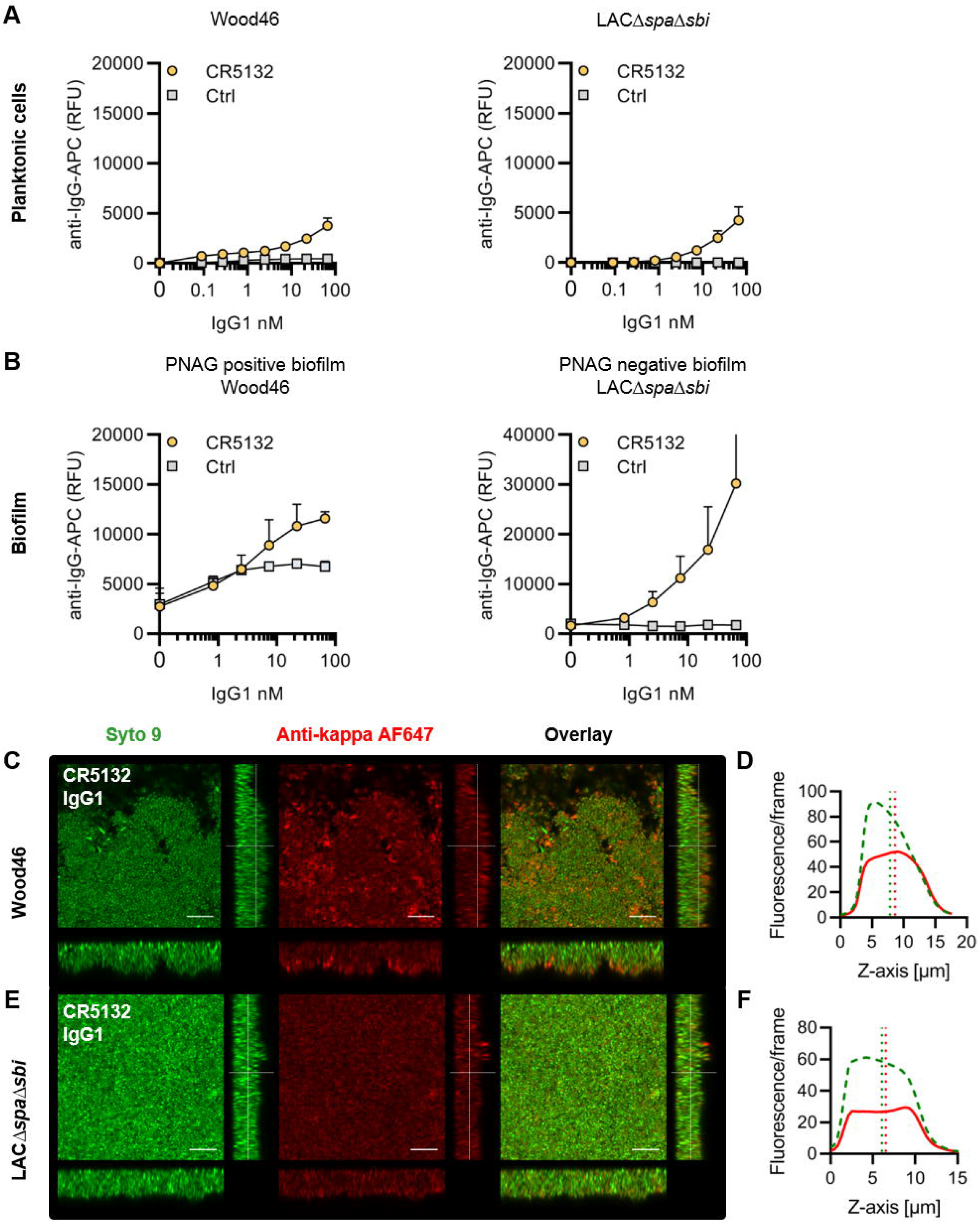
CR5132-IgG1 discriminates between planktonic bacteria and biofilm. (A) Planktonic bacteria of Wood46 (left) and LACΔ*spa*Δ*sbi* (right) were grown to exponential phase and incubated with a concentration range of CR5132-IgG1. MAb binding was detected using APC-labeled anti-human IgG antibodies and flow cytometry and plotted as geoMFI of the bacterial population. (B) Biofilm of Wood46 (left) and LACΔ*spa*Δ*sbi* (right) were grown for 24 h and incubated with a concentration range of CR5132-IgG1. MAb binding was detected using APC-labeled anti-human IgG antibodies and a plate reader and plotted as fluorescence intensity per well. Data represent mean + SD of at least 3 independent experiments. (C,D,E,F) Biofilm was grown for 24 h and incubated with 66 nM IgG1 mAb. Bacteria were visualized by Syto9 (green) and mAb binding was detected by staining with Alexa Fluor 647 conjugated goat-anti-human-kappa F(ab’)2 antibody (red). (C,E) Orthogonal views are representative for a total of three Z-stacks per condition and at least 2 independent experiments. Scale bars: 10 μm. (D,F) Z-stack profile plotting the total fluorescence of Syto9 (green, dotted line) and AF647 (red line) per frame versus the depth (μm) of the corresponding Z-stack. Vertical green and red lines represent the center of mass of the total fluorescent signal.

### RF1-IgG1 against the SDR protein family binds S. aureus in planktonic and biofilm form

Finally, we tested whether mAbs recognizing proteins on the staphylococcal cell surface are able to bind *S. aureus* biofilm. MAb rF1 recognizes the SDR family of proteins, which is characterized by a large stretch of serine-aspartate dipeptide repeats (SDR) and includes *S. aureus* clumping factor A (ClfA), clumping factor B (ClfB), and SDR proteins C, D and E and three additional SDR-proteins from *Staphylococcus epidermidis* (65). Mab rF1 recognizes glycosylated SDR repeats that are present in all members of this protein family. Additionally, the well described mAb T1-2 recognizes SDR family member clumping factor A (ClfA) (66, 67). We confirmed effective binding of rF1-IgG1 to planktonic exponential cultures of Wood46 and LACΔ*spa*Δ*sbi* **(Fig. 4a)**. In addition, both PNAG-positive and PNAG-negative biofilm formed by these strains were bound by rF1-IgG1 **(Fig. 4b)**. T1-2-IgG1 binding to planktonic bacteria was only detectable in stationary LACΔ*spa*Δ*sbi* cultures **(Fig. S6)** and not in exponential cultures **(Fig. 4a)** because ClfA is known to be expressed in the stationary phase (68). Furthermore, effective binding of T1-2-IgG1 to LACΔ*spa*Δ*sbi* PNAG-negative biofilm was detected **(Fig. 4b)**. In contrast, we could not detect T1-2-IgG1 binding to Wood46 PNAG-positive biofilm **(Fig. 4b)**, suggesting a greater abundance of ClfA in PNAG-negative biofilm than PNAG-positive biofilm. Alternatively, PNAG could shield ClfA from T1-2-IgG1 binding. In conclusion, we show that rF1-IgG1 and T1-2-IgG1 bind surface proteins on planktonic bacteria as well as biofilm formed by these bacteria. This means that besides *S. aureus* surface polysaccharides, surface proteins in a biofilm can also be recognized by mAbs.

**Figure 4.**
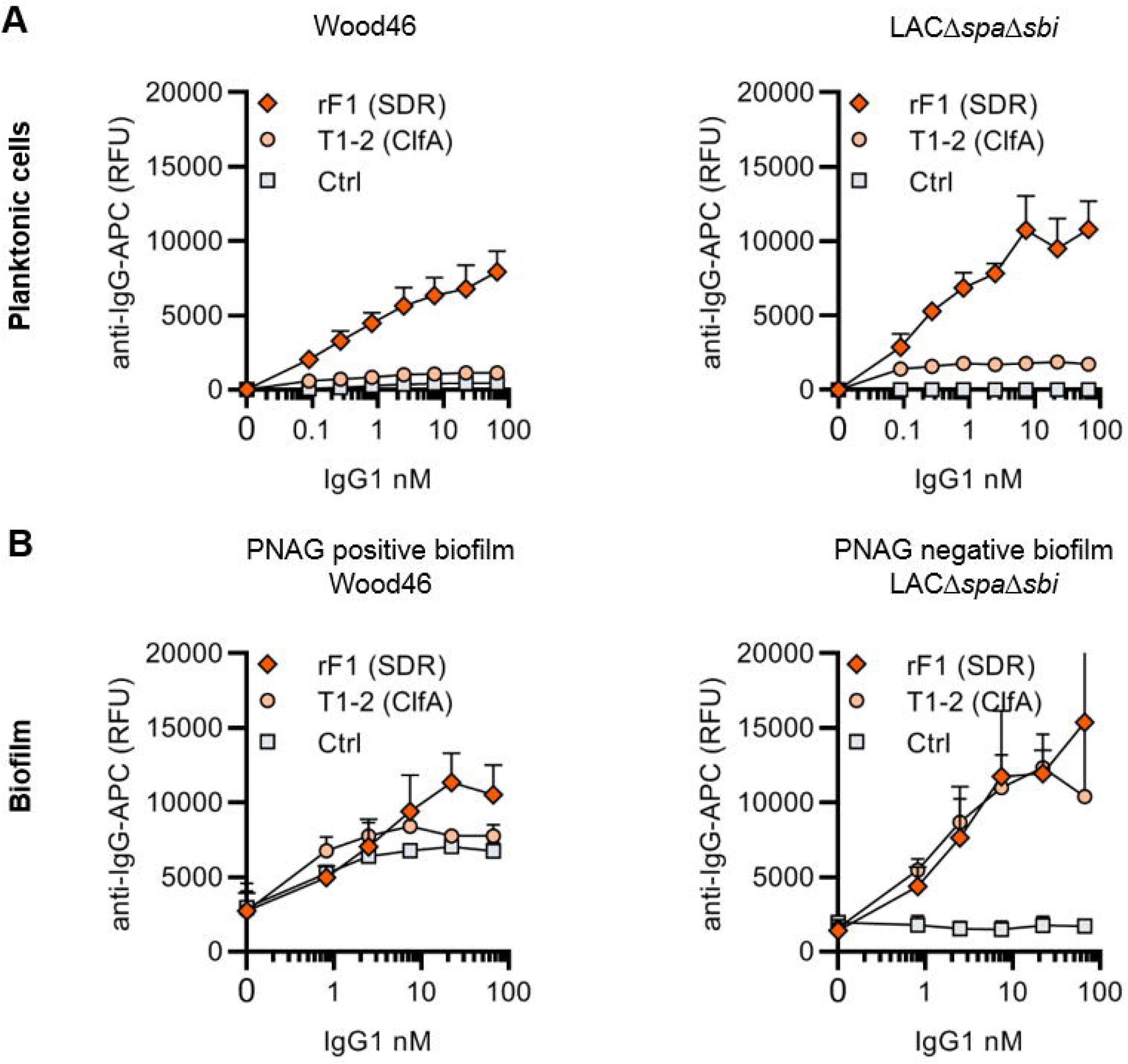
IgG1 mAbs against protein components bind planktonic bacteria as well as biofilm. (A) Planktonic bacteria of Wood46 (left) and LACΔ*spa*Δ*sbi* (right) were grown to exponential phase and incubated with a concentration range of rF1-IgG1 or T1-2-IgG1. MAb binding was detected using APC-labeled anti-human IgG antibodies and flow cytometry and plotted as geoMFI of the bacterial population. (B) Biofilm of Wood46 (left) and LACΔ*spa*Δ*sbi* (right) were grown for 24 h and incubated with a concentration range of rF1-IgG1 or T1-2-IgG1. MAb binding was detected using APC-labeled anti-human IgG antibodies and a plate reader and plotted as fluorescence intensity per well. Data represent mean + SD of at least 3 independent experiments.

### Comparative binding of mAbs to S. aureus biofilm

A direct comparison of all biofilm-binding mAbs revealed 4497-IgG1 as the best binder to PNAG-positive biofilm and CR5132 as the best binder to PNAG-negative biofilm **(Fig. 5ab)**. Furthermore, all mAbs that bind to planktonic bacteria **(Fig. S7)** were able to bind biofilm **(Fig. 5ab)** formed by that strain. Additionally, some mAbs, *i.e.* F598-IgG1 (anti-PNAG) and CR5132-IgG1 (anti-β-GlcNAc WTA), showed enhanced binding to biofilm compared to planktonic bacteria. Thus, we can identify two classes of monoclonal antibodies: one class recognizing both planktonic bacteria and biofilm, and one class recognizing biofilm only **(Table 1)**. Importantly, the mean AF647 fluorescence levels of Z-stacks acquired with the microscope corresponded to our plate reader data **(Fig. 5, S8)**. As most humans possess antibodies against *S. aureus*, we wondered whether pre-existing antibodies might compete with the IgG1 mAbs for binding to epitopes. To test this possibility, biofilm cultures were incubated with AF647-labeled mAbs in the presence of excess IgG (mAb:IgG ratio 1:25) isolated from pooled human serum. Despite the excess IgG, the AF647-labeled mAbs retained, on average, approximately 60% of the fluorescence they had in the absence of IgG **(Fig. S9)**. This indicates that the mAbs are able to recognize *S. aureus* biofilm in the presence of pre-existing antibodies.

**Figure 5.**
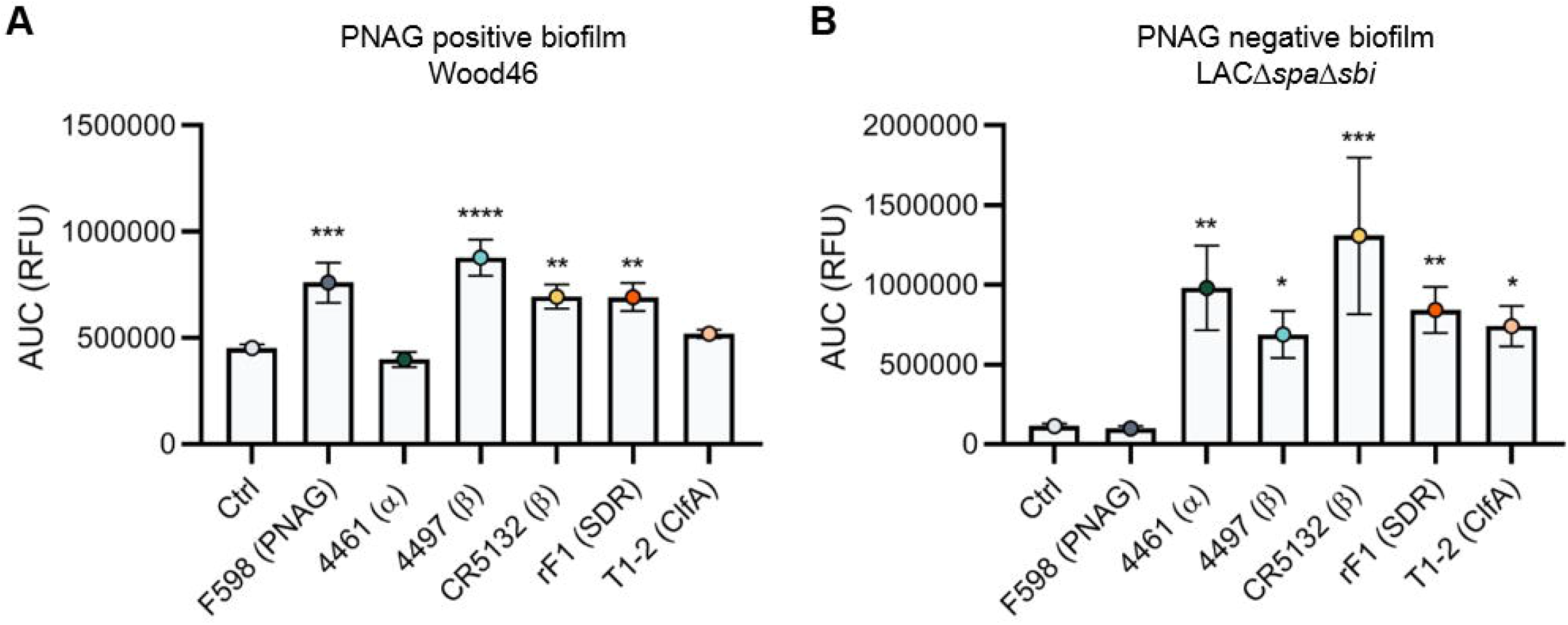
Comparative binding of IgG1 mAbs to *S. aureus* biofilm. Biofilm of Wood46 (A) and LACΔ*spa*Δ*sbi* (B) were grown for 24 h and incubated with a concentration range of IgG1 mAbs. MAb binding was detected using APC-labeled anti-human IgG antibodies and a plate reader. Data (A) were expressed as AUC of the binding curve (mean + SD) of at least 3 independent experiments. One-way ANOVA followed by Dunnett test was performed to test for differences in antibody binding versus control and displayed only when significant as *P ≤ 0.05, **P ≤ 0.01, ***P ≤ 0.001, or ****P ≤ 0.0001.

**Table 1.** MAb binding to biofilm and planktonic bacteria. Binding of mAbs is indicated with “+” and no binding is indicated with “–“.

| Clone | Target | Biofilm |  | Planktonic |  |
| --- | --- | --- | --- | --- | --- |
| | | PNAG (+) | PNAG (-) | Wood46 | LAC $\Delta spa\Delta sbi$ |
| F598 | PNAG | + | - | - | - |
| 4461 | WTA($\alpha$) | - | + | - | + |
| 4497 | WTA($\beta$) | + | + | + | - |
| CR5132 | WTA($\beta$) | + | + | - | +/- |
| rF1 | SDR proteins | + | + | + | + |
| T1-2 | ClfA | - | + | - | - |

### Indium-111 labelled 4497-IgG1 localizes to subcutaneous implant infections in an in vivo mouse model

Lastly, we studied whether mAbs against *S. aureus* biofilm could be used to localize a subcutaneous implant infection *in vivo*. Mice received a 5 mm catheter that was pre-colonized with *S. aureus* biofilm in one flank. As control, a sterile catheter was inserted into the other flank. Pre-colonized catheters were generated by incubating catheters with *S. aureus* USA300 LAC (WT) for 48 h. Bacterial loads on the catheters before implantation were approximately 4.5 × 10^7^ CFU **(Fig. S10a)**. We selected 4497-IgG1 (against β-GlcNAc WTA) because it potently binds to LAC biofilm *in vitro* **(Fig 5)**. To detect antibody localization in the mouse model, we radiolabelled 4497-IgG1 and control IgG1 with indium-111 (^111^In). Two days after implantation of the catheters, mice were injected intravenously with ^111^In-labeled 4497-IgG1 or ctrl-IgG1 and distribution of the radiolabel was visualized with total-body SPECT-CT scans at 24, 72 and 120 h after injection. Maximum intensity projections of SPECT/CT scans showed typical distribution patterns for IgG distribution in mice for both 4497 and ctrl antibodies (69, 70). At 24 h, activity was detected in blood-rich organs such as heart lungs and liver **(Fig. 6a, S11, movie S1,S2)**. Over time, antibodies were cleared from the circulation and blood-rich organs while the specific activity of radiolabeled 4497-IgG1 around pre-colonized implants remained. Remaining activity that was detected at incision sites of the pre-colonized catheters following both 4497-IgG1 and ctrl-IgG1 injection was likely explained by nonspecific accumulation of antibodies at inflammatory sites.

**Figure 6.**
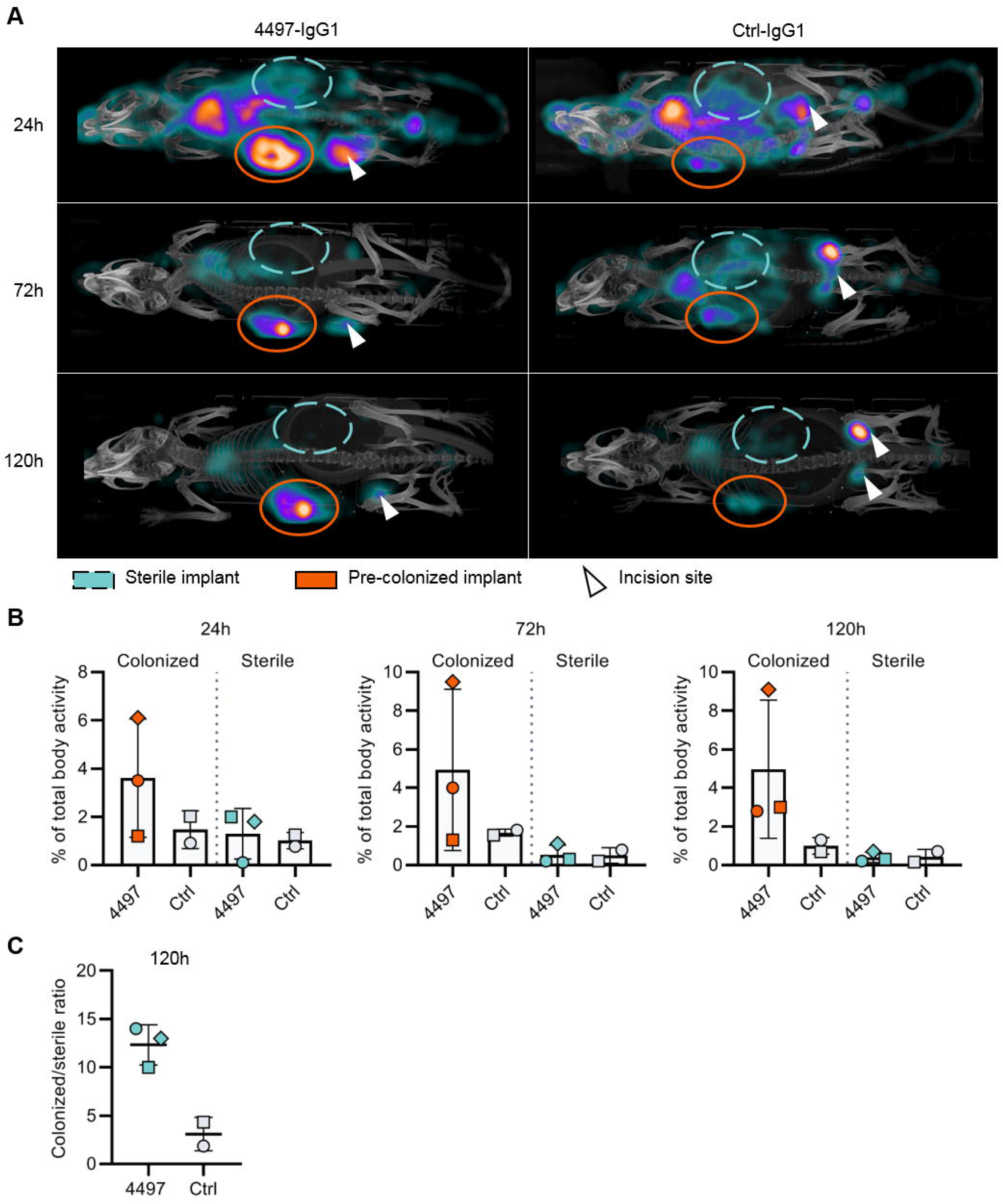
Localization of [^111^In]In-4497-IgG1 to a subcutaneous implant infection. (A) Maximum intensity projections (corrected for decay) of [^111^In]In-4497-IgG1 or [^111^In]In-Palivizumab injected in mice subcutaneously bearing pre-colonized (orange, full ellipses) and sterile (blue, striped ellipses) catheters. The mice were imaged at 24 h, 72 h and 120 h. Additional scans can be seen in the supplementary information. (B) The activity detected in regions of interests as indicated in (A) and supplementary information was expressed as percentage of total body activity. (C) For each individual mouse, percentages of total body activity around colonized implants were divided by activity around sterile implants.

To quantify the amount of antibody accumulating at pre-colonized and sterile implants, a volume of interest was drawn manually around the implants visible on SPECT-CT. The activity measured in the volume of interest was quantified as a percentage of the total body activity **(Fig. 6b)**. At all time points, 4497-IgG1 accumulated selectively at the pre-colonized catheter with a mean of 3.5% (24 h), 4.9% (72 h) and 5.0% (120 h) of the total body activity in the region of interest around the pre-colonized implant compared to 0.4% (24 h), 0.5% (72 h) and 0.4% (120 h) around the sterile implant. Ctrl-IgG1 was detected at pre-colonized catheters with a mean of 1.5% (24 h), 1.7% (72 h) and 0.7% (120 h) of the total body radiolabel activity and at sterile catheters with 0.8% (24 h), 0.2% (72 h) and 0.2% (120 h). Although there was no statistical difference in localization to pre-colonized implants between 4497-IgG1 and ctrl-IgG1, the colonized to sterile ratio at 120 h of 4497-IgG1 (mean ratio 12.3) was 4-fold higher than that for ctrl-IgG1 (mean ratio 3.1) **(Fig. 6C)**. Furthermore, the same results were found in a similar pilot experiment with less mAbs administered **(Fig. S12)**. At time point 120 h, thus 5 days after implantation of the catheter, a mean of ~1.1 × 10^6^ CFU were recovered from pre-colonized implants, whereas no bacteria were recovered from sterile controls, indicating that the infection had not disseminated to other locations in the body **(Fig. S10b)**. Interestingly, a higher bacterial burden recovered from pre-colonized implant correlated with a higher 4497-IgG1 activity at the implant **(Fig. 6a, S10b)**, indicating that a larger infection recruits more specific antibodies.

## Discussion

Although there is increasing interest in the use of monoclonal antibodies as therapeutic agents against *S. aureus* infection, only a few studies have focused on identifying antibodies that recognize its biofilm. Thus far, most efforts into mAb development against *S. aureus* have concentrated on antibodies that function by neutralization of *S. aureus* virulence factors. Identification of mAbs against *S. aureus* biofilms is a crucial starting point for the treatment and diagnosis of implant- or catheter-related infections. In this study we show that previously identified mAbs against *S. aureus* surface structures have the capacity to bind *S. aureus* biofilm. At the start of this study, the only mAb known to react with *S. aureus* biofilm was the F598 antibody recognizing PNAG. F598 was selected to bind to planktonic *S. aureus* MN8m, which is a spontaneous PIA/PNAG-overproducing mutant of strain Mn8 (71). Because numerous studies have shown that *S. aureus* is capable of forming different biofilm matrices (PNAG-positive and PNAG-negative) (4, 28, 29), we here focused on identifying antibodies recognizing different biofilm forms. Our study identified several mAbs **(Fig. 5)** capable of binding both types of biofilm (4497-, CR5132- and rF1-IgG1). This indicates that mAbs directed against WTA or the SDR protein family may be interesting candidates for treatment or diagnosis of *S. aureus* biofilm infections. WTA comprises ~30% of the *S. aureus* bacterial surface and therefore it is an attractive mAb target (57). However, WTA glycosylation can be strain specific and *S. aureus* can adapt WTA glycosylation upon environmental cues (58). Indeed, we found that 4461-IgG1 (anti-α-GlcNAc WTA) and 4497-IgG1 (anti-β-GlcNAc WTA) recognized different *S. aureus* strains and their biofilm. Thus, mAb therapy targeting WTA may best be composed of a mix of mAbs recognizing both α- and β-glycosylated WTA.

Our study also shows that it is possible for antibodies to recognize both *S. aureus* biofilm and planktonic bacteria. This is crucial because during biofilm infection, individual bacteria can disperse from the biofilm by secretion of various enzymes and surfactants to degrade the EPS (36). These dispersed bacteria can then disseminate and colonize new body sites or develop into sepsis, which is the most serious complication of biofilm-associated infections. With antibodies recognizing both biofilms and planktonic bacteria (like mAbs recognizing WTA (4461, 4497) and SDR protein family (rF1), it should be possible to target *S. aureus* bacteria *in vivo* throughout the entire infection cycle **(Table 1)**. We also observed that some mAbs (F598 and CR5132) bind better to biofilm than the planktonic form of *S. aureus*. Such antibodies might be useful for development of assays to discriminate between biofilm and planktonic cultures. Importantly, none of the mAbs in our panel bound planktonic *S. aureus* but not biofilm produced by the same strain. Our study should be taken as a promising proof-of-concept study in which we demonstrate binding of purified mAbs to two model *S. aureus* strains. As antigen expression is dependent on genetic variation between strains, future studies should extend these analyses for a large number of relevant strains and clinical isolates. Furthermore, mAb binding to *in vivo* biofilm could be complicated by the incorporation of host factors such as fibrinogen in the biofilm EPS (72–75).

Altogether, our *in vitro* data suggested that mAbs against *S. aureus* surface antigens may be suited to target biofilms *in vivo*. As a proof of principle, we tested ^111^In-labeled 4497-IgG1 localization in a mouse implant infection model and found increased radiolabel around the infected implant compared to the sterile implant within 24 h after mAb injection, suggesting rapid localization of 4497-IgG1 to biofilm *in vivo*. We consider these results as an excellent starting point to further evaluate the diagnostic and therapeutic purposes of these mAbs, not only for the treatment of *S. aureus* biofilms but also for other biofilm forming species. For advanced diagnostic purposes, specific mAbs could also be coupled to gamma- or positron emitting radionuclides and then be used to detect the presence of *S. aureus* in a biofilm in a patient or during revision surgery. Alternatively, mAbs could be used *in vitro* to detect the presence of biofilm on explanted implants. For therapeutic purposes, mAbs could function as a delivery vehicle to specifically direct biofilm degrading enzymes, antibiotics, photosensitizers or alpha/beta emitting radionuclides to the site of infection. Alternatively, mAbs that bind to biofilms could induce the activation of the immune system via the Fc-domain(5). Therapeutic mAbs could be used as adjunctive therapy in chronic infections or as a prophylactic treatment after replacement of an infected implant to prevent colonization of the new implant. In all cases, an effective antibody therapy against *S. aureus* biofilm will have vast utility in patients undergoing medical procedures.

## Materials and Methods

### Ethics statement

All animal procedures were approved by the Utrecht University animal ethics committee and were performed in accordance with international guidelines on handling laboratory animals (Animal Use Permit #AVD1150020174465, approved 1 March 2018). To obtain human serum, blood was isolated from healthy donors according to a study protocol approved by the Medical Ethics Committee of the University Medical Center Utrecht was obtained (METC protocol 07-125/C approved on March 1, 2010).

### Expression and isolation of human monoclonal antibodies

For human monoclonal antibody expression, variable heavy (VH) and light (VL) chain sequences were cloned in homemade pcDNA3.4 expression vectors containing human heavy chain (HC) and light chain (LC) constant regions, respectively. To generate these homemade HC and LC constant region expression vectors, HC and LC constant regions from pFUSE-CHIg-hG1 and pFUSE-CLIg-hk (Invivogen) were amplified by PCR and cloned separately into pcDNA3.4 (ThermoFisher Scientific). All sequences used are shown in **Table S1**. VH and VL sequences were derived from antibodies previously described in scientific publications and patents listed in **Table S1**. Originally, all antibodies have been described as fully human, except for T1-2 which was raised in mice by immunization with ClfA (76) and later humanized to T1-2 (67). CR5132 was discovered using ScFv phage libraries [US 2012/0141493 A1] and F598 (71), 4461, 4497 (38) and rF1 (40) were cloned from human B cells derived from *S. aureus* infected patients. For each VH and VL, human codon-optimized genes with an upstream KOZAK sequence and a HAVT20 signal peptide (MACPGFLWALVISTCLEFSMA) were ordered as gBlocks (Integrated DNA Technologies) and cloned into pcDNA3.4 HC and LC constant region expression vectors using Gibson assembly (Bioke). TOP10F’ *E. coli* were used for propagation of the generated plasmids. After sequence verification plasmids were isolated using NucleoBond Xtra Midi plasmid DNA purification (Macherey-Nagel). For recombinant antibody expression, 2×10^6^ cells/ml EXPI293F cells (Life Technologies) were transfected with 1 μg DNA/ml cells in a 3:2 (LC:HC) ratio were transfected using polyethylenimine HCl MAX (Polysciences). After 4 to 5 days of expression, IgG1 antibodies were isolated from cell supernatant using a HiTrap protein A column (GE Healthcare) and Äkta Pure protein chromatography system (GE Healthcare). Antibody fractions were dialyzed overnight in PBS and filter-sterilized though 0.22 μm Spin-X filters. Antibodies were analyzed by size exclusion chromatography (GE Healthcare) and separated for monomeric fraction in case aggregation levels were >5%. Antibody concentration was determined by measurement of the absorbance at 280 nm and stored at −20°C.

### Bacterial strains and growth conditions

*S. aureus* strains Wood46 (ATCC 10832) (46, 47, 77), USA300 LAC (AH1263) (29) and USA300 LAC Δ*spa, sbi*∷Tn (AH4116) were used in this study. Strain USA300 LAC Δ*spa, sbi*∷Tn (AH4116) was constructed by transducing *sbi*∷Tn from Nebraska Transposon Library (78) into USA300 LAC Δ*spa* (AH3052) (79) with phage 11. Strains were grown overnight on sheep blood agar (SBA) at 37°C and were cultured overnight in Tryptic Soy Broth (TSB) before each experiment. For exponential phase planktonic cultures, overnight cultures were sub-cultured in fresh TSB for 2 h. For stationary phase planktonic cultures, overnight cultures in TSB were used.

### Biofilm culture

For PNAG-negative biofilm, overnight cultures of LAC or LAC Δ*spa sbi*∷Tn were diluted to an OD_600_ of 1 and then diluted 1:1000 in fresh TSB containing 0.5% (wt/vol) glucose and 3% (wt/vol) NaCl. 200 μl was transferred to wells in a flat bottom 96 wells plate (Corning costar 3598, Tissue Culture treated) and incubated statically for 24 h at 37°C. To facilitate attachment of PNAG-negative bacteria to the wells, plates were coated overnight at 4°C before inoculation. For experiments with EPS degrading enzymes, plates were coated with 20% human plasma (Sigma) in carbonate – bicarbonate buffer. For IgG1 binding assays, plates were coated with 20 μg/ml human fibronectin (Sigma) in 0.1M carbonate – bicarbonate buffer (pH 9.6). PNAG-positive Wood46 biofilms were grown similarly, except that no coating was used and growth medium was TSB supplemented with 0.5% (wt/vol) glucose.

### Antibody binding to planktonic cultures

To determine mAb binding capacity, planktonic bacterial cultures were suspended and washed in PBS containing 0.1% BSA (Serva) and mixed with a concentration range of IgG1-mAbs in a round-bottom 96-well plate in PBS-BSA. Each well contained 2.5 × 10^6^ bacteria in a total volume of 55 μl. Samples were incubated for 30 min at 4°C, shaking (~700 rpm) and washed once with PBS-BSA. Samples were further incubated for another 30 min at 4°C, shaking (~700 rpm), with APC-conjugated polyclonal goat-anti-human IgG F(ab’)2 antibody (Jackson immunoresearch, 1:500). After washing, samples were fixed for 30 min with cold 1% paraformaldehyde. APC fluorescence per bacterium was measured on a flow cytometer (FACSVerse, BD). Control bacteria were used to set proper FSC and SSC gate definitions to exclude debris and aggregated bacteria. Data were analyzed with FlowJo (version 10).

### Antibody binding to biofilm cultures

To determine mAb binding capacity to biofilm, wells containing 24 h biofilm were blocked for 30 min with 4% BSA in PBS. After washing with PBS, wells were incubated with a concentration range of IgG1-mAbs, or Fab fragments when indicated, in PBS-BSA (1%) for 1 h at 4°C, statically. After washing two times with PBS, samples were further statically incubated for 1 h at 4°C with APC-conjugated polyclonal goat-anti-human IgG F(ab’)2 antibody (Jackson immunoresearch, 1:500). Fab fragments were detected with Alexa Fluor 647 conjugated goat-anti-human-kappa F(ab’)2 antibody (Southern Biotech, 1:500). After washing, fluorescence per well was measured using a CLARIOstar plate reader (BMG LABTECH).

### Confocal microscopy of static biofilm

Wood46 and LAC Δ*spa sbi∷*Tn biofilm were grown in glass bottom Cellview™ slides (Greiner bio-one (543079)) similarly as described above. Cellview™ slides were placed in a humid chamber during incubation to prevent evaporation of growth medium. After 24 h, wells were gently washed with PBS and fixed for 30 min with cold 1% paraformaldehyde, followed by blocking with 4% BSA in PBS. After washing with PBS, wells were incubated with 66 nM IgG1-mAbs in PBS-BSA (1%) for 1 h at 4°C, statically. After washing two times with PBS, samples were further statically incubated for 1 h at 4°C with Alexa Fluor 647 conjugated goat-anti-human-kappa F(ab’)_2_ antibody (Southern Biotech, 1:300) and 6 μM Syto9 (Live/Dead BacLight Bacterial Viability Kit; Invitrogen). Z-stacks at 3 random locations per sample were collected at 0.42 μm intervals using a Leica SP5 confocal microscope with a HCX PL APO CS 63×/1.40–0.60 OIL objective (Leica Microsystems). Syto9 fluorescence was detected by excitation at 488 nm and emission was collected between 495nm and 570nm. Alexa Fluor 647 fluorescence was detected by excitation at 633 nm and emission was collected between 645 nm - 720 nm. Image acquisition and processing was performed using Leica LAS AF imaging software (Leica Microsystems).

### Subcutaneous implant infection mouse model

To determine *in vivo* mAb localization to implant associated infections, a subcutaneous implant infection mouse model was developed based on Kadurugamuwa *et al*.(80). Five Balb/cAnNCrl male mice weighing >20 g obtained from Charles River Laboratories were housed in our Laboratory Animal Facility. One hour before surgery, all mice were given 5 mg/kg Carprofen. Anesthesia was induced with 5% isoflurane and maintained with 2% isoflurane. Their backs were shaved and the skin was disinfected with 70% ethanol. A 5 mm skin incision was made using scissors after which an 14 Gauge piercing needle was carefully inserted subcutaneously at a distance of approximately 1-2 cm. A 5 mm segment of a 7 French polyurethane catheter (Access Technologies) was inserted into the piercing needle and correctly positioned using a k-wire. The incision was closed using one or two sutures and the skin was disinfected with 70% ethanol. Mice received one s.c. catheter in each flank. One catheter served as a sterile control whereas the other was pre-colonized for 48 hours with an inoculum of ~10^7^ CFU *S. aureus* LAC. Before inoculation, the implants were sterilized with 70% ethanol and air dried. The inoculated implants were incubated at 37°C for 48 h under agitation (200-300 RPM). New growth medium was added at 24 h to maintain optimal growing conditions. Implants were washed three times with PBS to remove non-adherent bacteria and stored in PBS until implantation or used for determination of viable CFU counts. To this end, implants were placed in PBS and sonicated for 10 min in a Branson M2800E Ultrasonic Waterbath (Branson Ultrasonic Corporation). After sonication, total viable bacterial counts per implant were determined by serial dilution and plating.

### Radionuclides and radiolabeling of antibodies

4497-IgG1 (anti-β-GlcNAc WTA) and control IgG1 antibody Palivizumab (MedImmune) were labeled with indium-111 (^111^In) using the bifunctional chelator CHXA” as described previously by Allen et al (69). In short, antibodies were buffer exchanged into conjugation buffer and incubated at 37°C for 1.5 h with a 5-fold molar excess of bifunctional CHXA” (Macrocyclics, prepared less than 24 h before use). The mAb-CHXA” conjugate was then exchanged into 0.15 M ammonium acetate buffer to remove unbound CHXA” and subsequently incubated with approximately 150 kBq ^111^In (purchased as [^111^In]InCl_3_ from CuriumPharma) per μg mAb. The reaction mixture was incubated for 60 min at 37°C after which free ^111^In^3+^ was quenched by the addition of 0.05 M EDTA. Quality control was done by instant thin layer chromatography (iTLC) and confirmed radiolabeling at least 95 % radiochemical purity of the antibodies.

### USPECT-CT and CFU count

Two days after subcutaneous implantation of catheters, 50 μg radiolabeled antibody (7.5 MBq) was injected into the tail vein. Three mice were injected with [^111^In]In-4497-IgG1 and 2 mice were injected with [^111^In]In-Palivizumab. At 24, 72 and 120 h post injection, multimodality SPECT/CT imaging of mice was performed with a VECTor^6^ CT scanner (MILabs, The Netherlands) using a MILabs HE-UHR-M mouse collimator with 162 pinholes (diameter, 0.75 mm) (81). At 24 h, a 30 min total-body SPECT-CT scan was conducted under anesthesia. Scanning duration at 72 and 120 h was corrected for the decay of ^111^In. Immediately after the last scan, mice were sacrificed by cervical dislocation while under anesthetics. The carcasses were stored at −20 °C until radiation exposure levels were safe for further processing. Implants were aseptically removed, placed in PBS and sonicated for 10 min in a Branson M2800E Ultrasonic Waterbath (Branson Ultrasonic Corporation). After sonication, total viable bacterial counts per implant were determined by serial dilution and plating. SPECT image reconstruction was performed using Similarity Regulated OSEM (82), using 6 iterations and 128 subsets. Image processing and volume of interest analysis of the total-body SPECT scans were done using PMOD (PMOD Technologies). The total-body SPECT volumes were smoothed using a 3D gaussian filter of 1.5 mm. To quantify the accumulation of ^111^In around the implants, corresponding 3D regions of interest were manually drawn around catheters that were located in SPECT-CT fusion scans. Iso-contouring of these specific regions were done using a threshold of 0.11 after which the voxel intensity sums were related to dose calibrator measurements.

### Statistical testing

Statistical analyses were performed in GraphPad Prism 8. The tests and n-values used to calculate p-values are indicated in the figure legends. Unless stated otherwise, graphs are comprised of at least three biological replicates.

## Acknowledgments

The authors greatly thank Reindert Nijland (Department of Animal Sciences, Wageningen University and Research, Wageningen, The Netherlands) and Fernanda Paganelli (Department of Medical Microbiology, UMC Utrecht) for assistance with biofilm work; Astrid Hendriks (Department of Medical Microbiology, UMC Utrecht), Nina van Sorge (Department of Medical Microbiology and Infection Prevention, Amsterdam UMC, The Netherlands) and Jeroen Codee (Leiden Institute of Chemistry, Leiden University, The Netherlands,) for providing WTA glycosylated beads; Frank Beurskens (Genmab B.V., Utrecht, The Netherlands) for help with selection of mAbs; Sonja von Aulock and Siegfried Morath (Department of Biochemical Pharmacology, University of Konstanz, Konstanz, Germany) for providing LTA preparates. L.d.V. and B.v.D. are supported by a grant from Health~Holland (LSHM17026 to J.v.S. and H.W.). F.J.B. and R.M.R are supported by the research grant QUARAT: Quantitative Universal Radiotracer Tomography (TTW16885, Dutch Research Council (NWO). A.R.H. was supported by a merit award (BX002711) from the U.S. Department of Veteran Affairs and grant AI083211 from the National Institutes of Health.

## Notes

### Competing Interest Statement

The authors have declared no competing interest.

