## Supplemental Methods, Figures and Tables for "Human monoclonal antibodies against *Staphylococcus aureus* surface antigens recognize *in vitro* biofilm and *in vivo* implant infections"

**This PDF file includes:**

Supplementary methods
Figures S1 to S12
Table S1
Legends for Movie S1 and Movie S2
SI References

**Other supplementary materials for this manuscript include the following:**

Movie S1
Movie S2

**Supplementary Methods**

*Crystal violet assay*

To determine the sensitivity of biofilms to DNase I, 1 mg/mL bovine DNase I (Roche) was added at the same time as inoculation and incubated during biofilm formation for 24 h. To determine biofilm sensitivity to DspB, 30 nM DspB (MTA-Kane Biotech Inc.) was added to 24 h biofilm and incubated statically for 2 h at 37°C. Biofilm adherence after treatment with DNase I or DspB compared to untreated controls was analyzed as follows. Wells were washed once with PBS and adherent cells where fixed by drying plates at 60°C for 1 h. Adherent material was stained with 0.1% crystal violet for 5 min and excess stain was removed by washing with distilled water. Remaining dye was solubilized in 33% acetic acid and biofilm formation was quantified by measuring the absorbance at 595 nm using a CLARIOstar plate reader (BMG LABTECH).

*Scanning electron microscopy*

Biofilms were grown as described above but on 12 mm round poly-L-Lysin coated glass coverslip (Corning). Coverslips were washed 1x with PBS and fixed for 24 h at room temperature with 2% (v/v) formaldehyde, 0.5% (v/v) glutaraldehyde and 0.15% (w/v) Ruthenium Red in 0.1 M Phosphate buffer (pH 7.4). Coverslips were then rinsed two times with phosphate buffer and post fixed for 2 h at 4°C with 1% osmium tetroxide and 1.5% (w/v), potassium ferricyanide (K3[Fe(CN)6]) in 0.065 M phosphate buffer (pH 7.4). Coverslips were rinsed once in distilled water followed by a stepwise dehydration with ethanol (i.e. 50%, 70%, 80%, 95%, 2x100%). Samples were then treated stepwise with hexamethyldisilizane (i.e. 50% HMDS/ethanol, 2X100% HMDS) and air-dried overnight. The next day samples were mounted on 12 mm aluminum stubs for SEM using carbon adhesive discs (Agar Scientific), additional conductive carbon tape (Agar Scientific) was placed over part of the sample to establish a conductive path to reduce charging effects. To further improve conductivity, the surface of the samples were coated with a 6 nm layer of Au using a Quorum Q150R S sputter coater. Samples were imaged with a Scios FIB-SEM (Thermo Scientific) under high-vacuum conditions, at an acceleration voltage of 20 kV and a current of 0.40 nA.

*Peptidoglycan and LTA ELISA*

Peptidoglycan from Wood46 was isolated as described in Timmerman *et al.*(1) and purified LTA was a kind gift from Sonja von Aulock and Siegfried Morath (University of Konstanz). We coated Maxisorb plates (Nunc) overnight at 4°C with 1 µg/ml peptidoglycan or LTA. The plates were washed three times with PBS 0.05% Tween, blocked with PBS 4% BSA, and incubated one hour with a concentration range of CR5132-IgG1, A120-IgG1 (directed against LTA) or control IgG1. The plates were washed, incubated 1 h with 1:6000 Goat-fab'2-anti-human-kappa-HRP (Southern Biotech). Finally, the plates were washed and developed using 3,3',5,5'-tetramethylbenzidine

(Thermo Fisher). The reaction was stopped by addition of 1 N $H_2SO_4$. Absorption at 450nm was measured using a CLARIOstar plate reader (BMG LABTECH).

*IgG1 binding to WTA glycosylated beads*

Synthetic WTA (a kind gift of Jeroen Codee, Leiden University) was immobilized on magnetic beads as in van Dalen *et al*.(2). Shortly, biotinylated RboP hexamers were enzymatically glycosylated by recombinant TarM, TarS or TarP with UDP-GlcNAc (Merck) as substrate. After 2 h incubation at room temperature, $5x10^7$ pre-washed Dynabeads M280 Streptavidin (Thermo Fisher) were added and incubated 15 min at room temperature. The coated beads were washed three times in PBS using a plate magnet, resuspended in PBS 0.1% BSA and stored at 4°C. To determine CR5132 binding capacity, beads were suspended and washed in PBS/0.05% Tween/0.1% BSA and mixed with a concentration range of CR5132-IgG1 or control IgG1 in a round-bottom 96-well plate in PBS/Tween/BSA. Each well contained $10^5$ beads. Samples were incubated for 30 min at 4°C, shaking (~700 rpm) and washed once with PBS/Tween/BSA. Samples were further incubated for another 30 min at 4°C, shaking (~700 rpm), with APC-conjugated polyclonal goat-anti-human IgG $F(ab')_2$ antibody (Jackson immunoresearch, 1:500). After washing, APC fluorescence per bead was measured on a flow cytometer (FACSVerse, BD).

*Antibody binding in the presence of human pooled IgG*

MAb binding in the presence of human pooled IgG was assessed with mAbs that were directly fluorescently-labeled. Briefly, mAbs were labeled with AF647 NHS ester (ThermoFisher Scientific) by following the manufacturer's protocol. Labeled mAbs were buffer exchanged into PBS using desalting Zeba columns (ThermoFisher Scientific), checked for degree of labeling (ranging from 2.9 to 4.5) and stored at 4°C. To isolate human pooled IgG, blood was drawn from 22 healthy volunteers and allowed to clot for 15 min at room temperature. After centrifugation for 10 minutes at 3,220 x g at 4°C, serum was collected, pooled and subsequently stored at -80°C. IgG was isolated from pooled serum as described above. Biofilm cultures were prepared, washed and incubated as described above. Samples were incubated with 10 µg/mL AF647-conjugated IgG1 mAbs in buffer or buffer containing 250 µg/mL pooled IgG. AF647 fluorescence per well was measured using a CLARIOstar plate reader (BMG LABTECH).

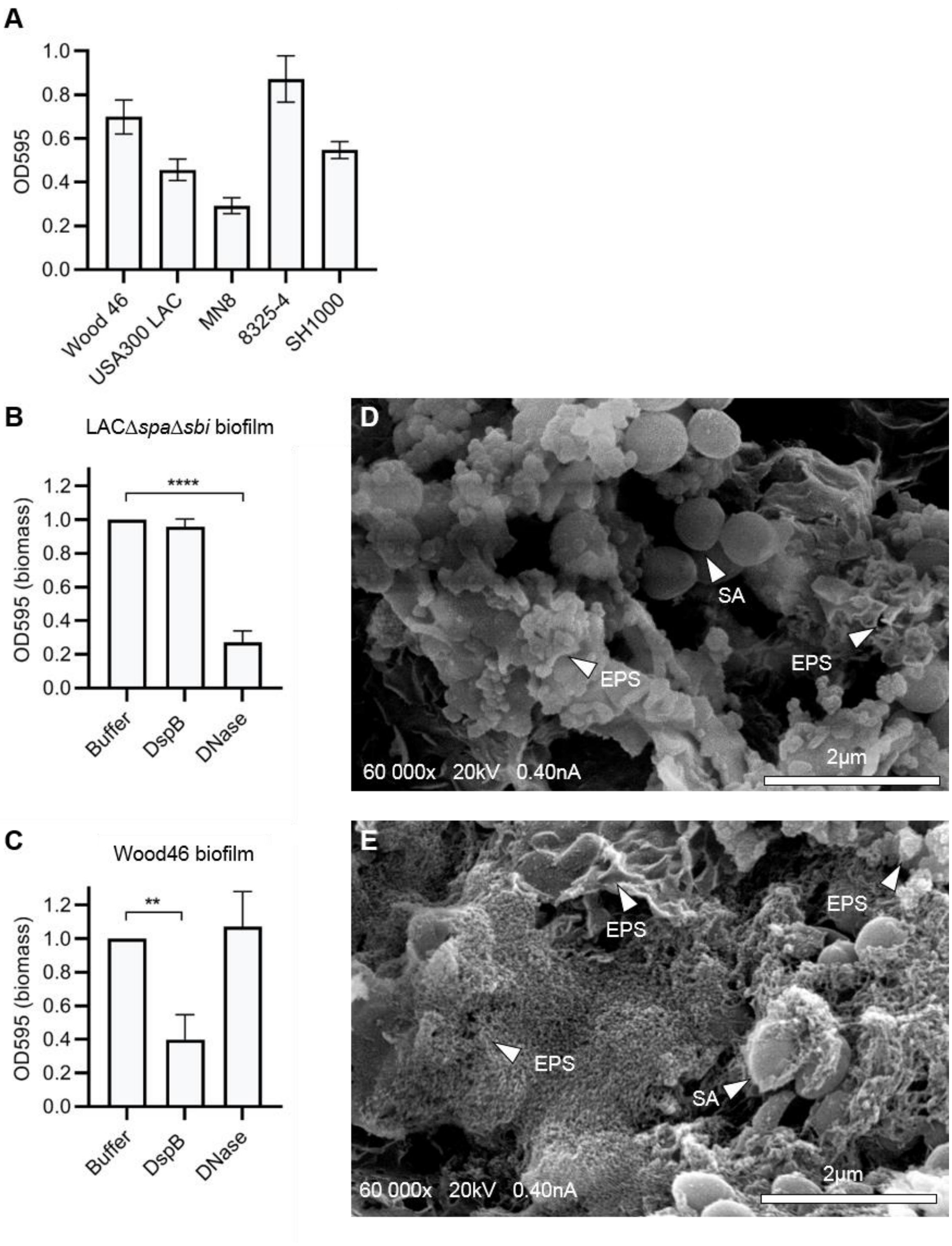


**Fig. S1.** *S. aureus* strains LAC and Wood46 form different types of biofilm. (A) Biofilm of different *S. aureus* strains available in our lab was grown for 24 h in TSB + 0.5% glucose on uncoated polystyrene 96-w plates. Adherent biofilm biomass was measured by crystal violet staining. Data represents mean + SD of 3 triplicates in 1 independent experiment. (B,D) Biofilm of *S. aureus* strain LACΔ*spa*Δ*sbi* (B) and Wood46 (D) was grown for 24 h with buffer or DNase (1 mg/mL). DspB (30 nM) was added after 24 h of biofilm formation. Adherent biofilm biomass was measured by crystal violet staining. (D,E) Representative SEM images of LAC∆*spa*∆*sbi* (D) and Wood 46 (E) biofilms established on glass coverslips following 24h incubation at 37°C. SA = *S. aureus*, EPS = Extracellular Polymeric Substance structure. Data represent mean + SD of 3 independent experiments. Triplicates were averaged and expressed as relative biomass by dividing the OD595 of treated samples by the OD595 of control samples. One-way ANOVA followed by Dunnett test was performed to test for differences in biofilm biomass and displayed only when significant as *P ≤ 0.05, **P ≤ 0.01, ***P ≤ 0.001, or ****P ≤ 0.0001.

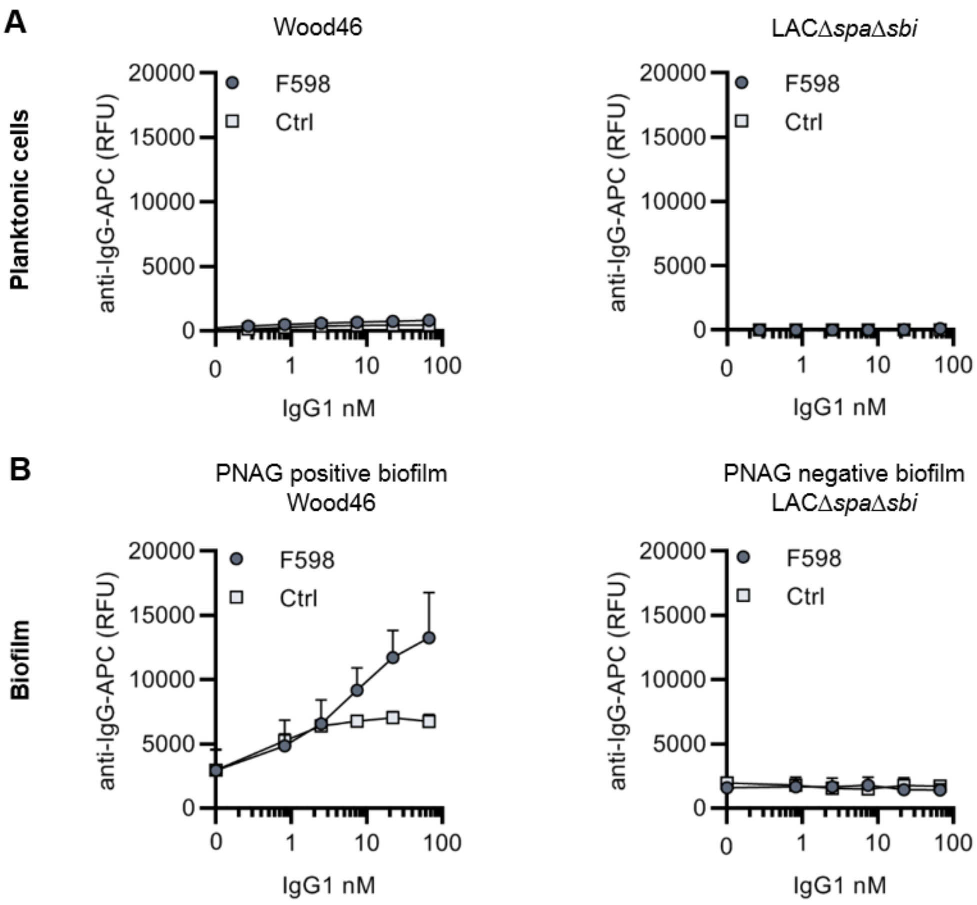


**Fig. S2.** F598-IgG1 binds PNAG-dependent biofilms specifically. (A) Planktonic bacteria of LACΔ*spa*Δ*sbi* (left) and Wood46 (right) were grown to exponential phase and incubated with a concentration range of F598-IgG1. MAb binding was detected using APC-labeled anti-human IgG antibodies and flow cytometry and plotted as geoMFI of the bacterial population. (B) Biofilm of Wood46 and LACΔ*spa*Δ*sbi* were grown for 24 h and incubated with a concentration range of F598-IgG1. MAb binding was detected using APC-labeled anti-human IgG antibodies and a plate reader and plotted as fluorescence intensity per well. Data represent mean + SD of at least 3 independent experiments.

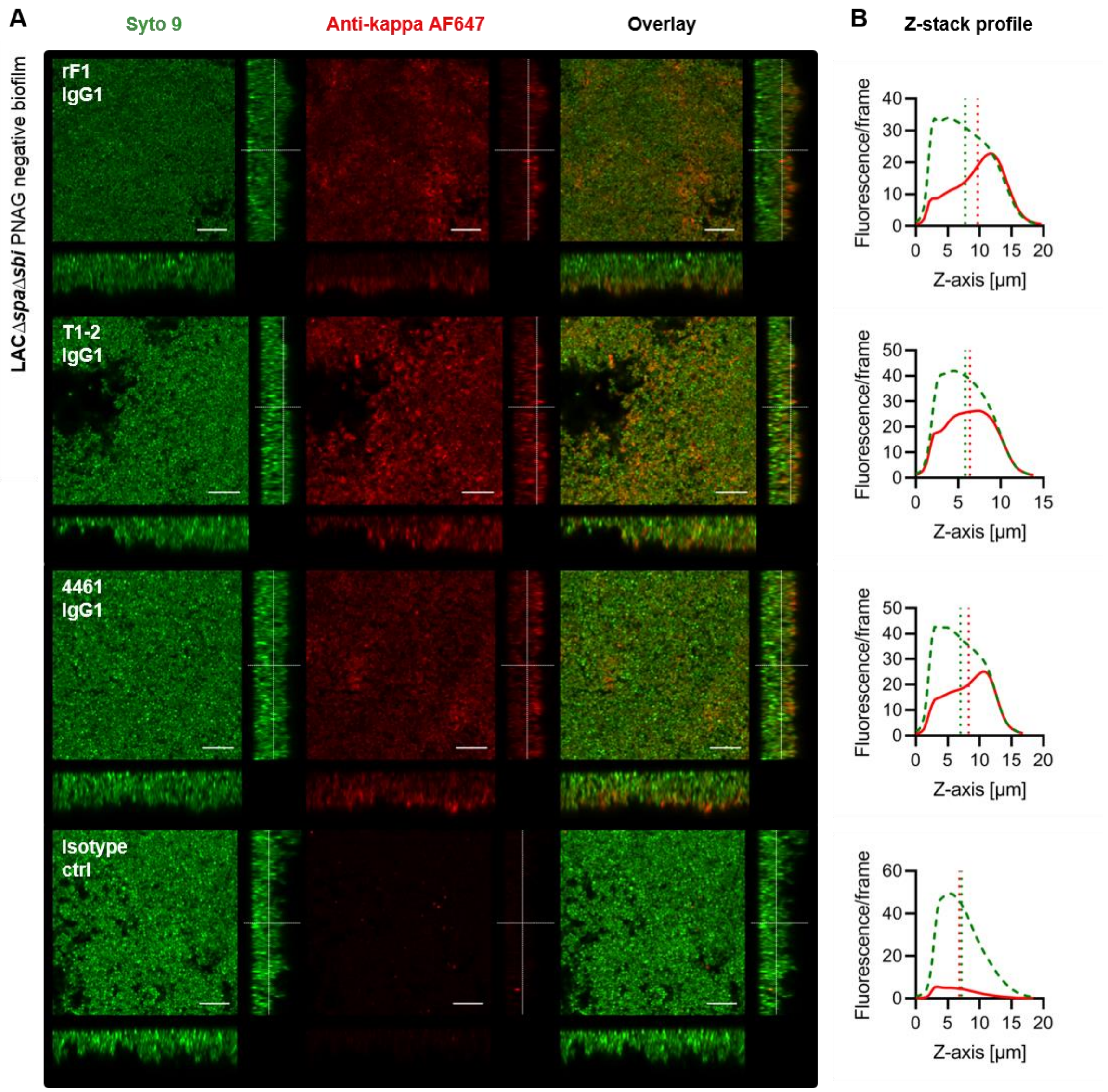


**Fig. S3.** Orthogonal views of PNAG-negative biofilm incubated with IgG1 mAbs. (A) Biofilm was grown for 24 h and incubated with 66 nM IgG1 mAbs or isotype controls. Bacteria were visualized by Syto 9 (green) and mAb binding was detected by staining with Alexa Fluor 647 conjugated goat-anti-human-kappa F(ab')$_2$ antibody (red). Syto 9 and AF647 were imaged using 488 and 633 nm lasers. Images are representative for a total of three Z-stacks per condition and at least 2 independent experiments. Scale bars: 10 µm. (B) Z-stack profile plotting the total fluorescence of Syto 9 (green, dotted line) and AF647 (red line) per frame versus the depth (µm) of the corresponding Z-stack. Vertical green and red lines represent the center of mass of the total fluorescent signal.

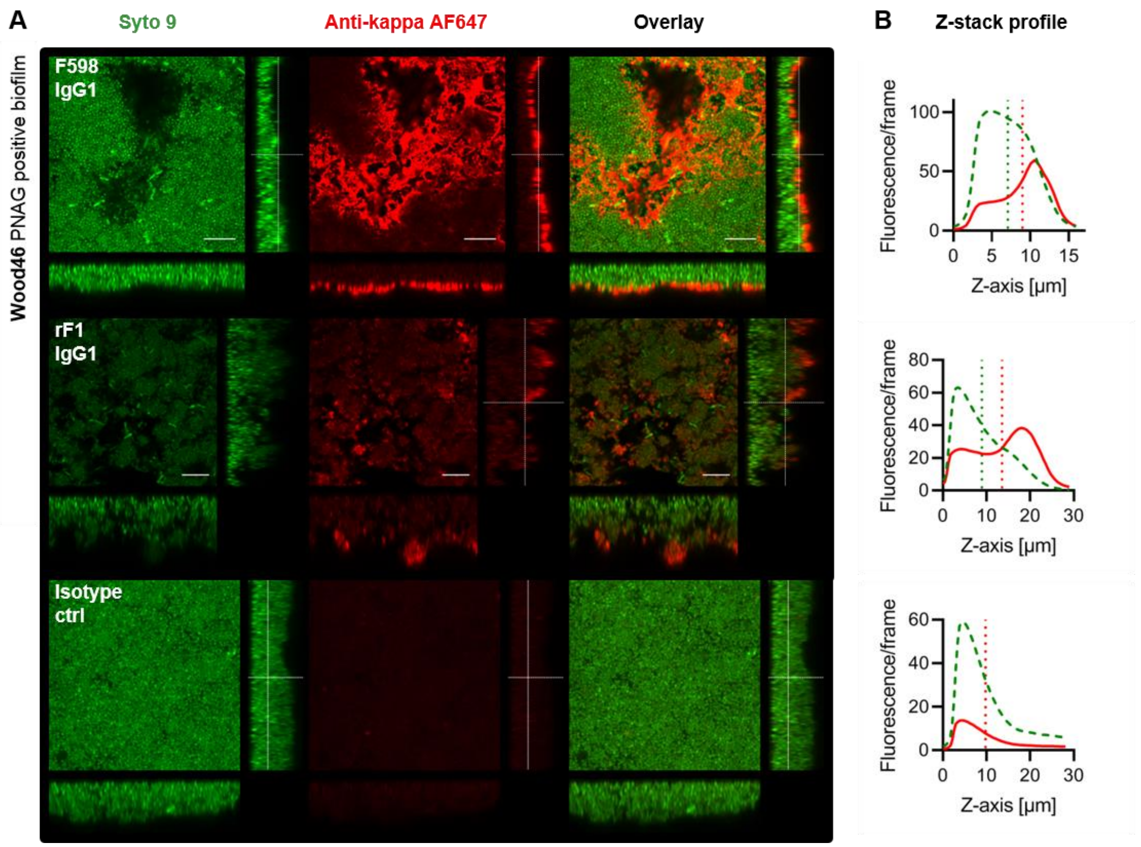


**Fig. S4.**Orthogonal views of PNAG-positive biofilm incubated with IgG1 mAbs. (A) Biofilm was grown for 24 h and incubated with 66 nM IgG1 mAbs or isotype controls. Bacteria were visualized by Syto 9 (green) and mAb binding was detected by staining with Alexa Fluor 647 conjugated goat-anti-human-kappa F(ab')$_2$ antibody (red). Syto 9 and AF647 were imaged using 488 and 633 nm lasers. Images are representative for a total of three Z-stacks per condition and at least 2 independent experiments. Scale bars: 10 µm. (B) Z-stack profile plotting the total fluorescence of Syto 9 (green, dotted line) and AF647 (red line) per frame versus the depth (µm) of the corresponding Z-stack. Vertical green and red lines represent the center of mass of the total fluorescent signal.

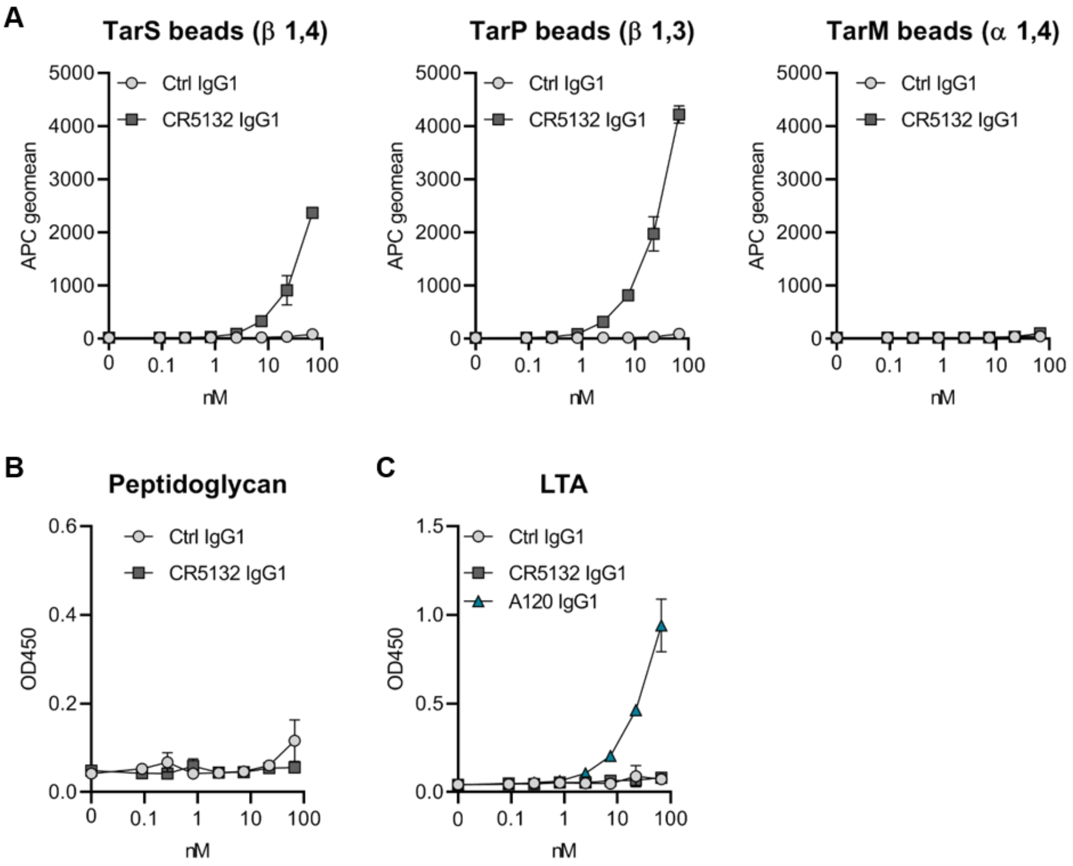


**Fig. S5.** Target identification of CR5132. (A) WTA coated beads were incubated with a concentration range of mAbs. MAb binding was detected using APC-labeled anti-human IgG antibodies and flow cytometry and plotted as geoMFI + SD of duplicates. (B, C) ELISA plates were coated with purified peptidoglycan (B) and LTA (C). Plates were incubated with a concentration range of mAbs and mAb binding was detected using anti-human kappa-HRP antibodies.

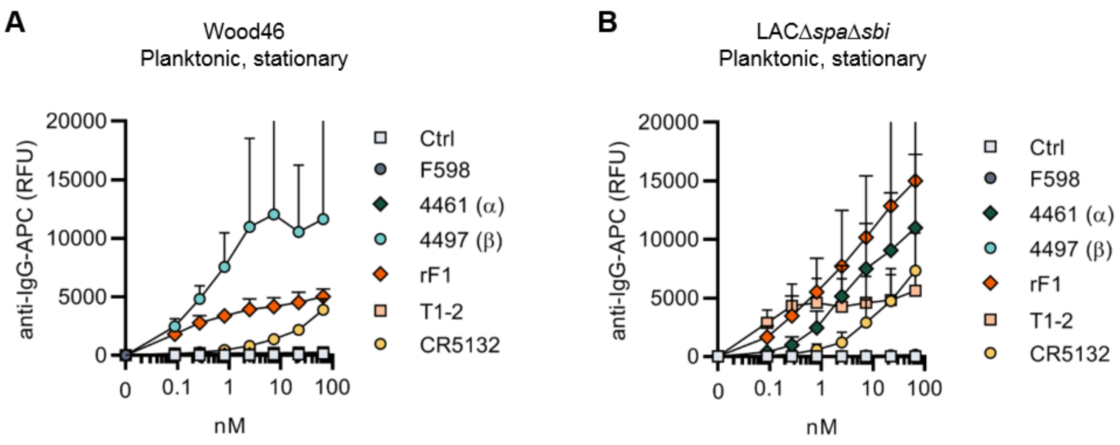


**Fig. S6.** Binding of the mAb panel to stationary phase planktonic cultures. Planktonic bacteria of Wood46 (A) LACΔ*spa*Δ*sbi* (B) and were grown to stationary phase and incubated with a concentration range of mAbs. MAb binding was detected using APC-labeled anti-human IgG antibodies and flow cytometry and plotted as geoMFI of the bacterial population. Data represent mean + SD of at least 3 independent experiments.

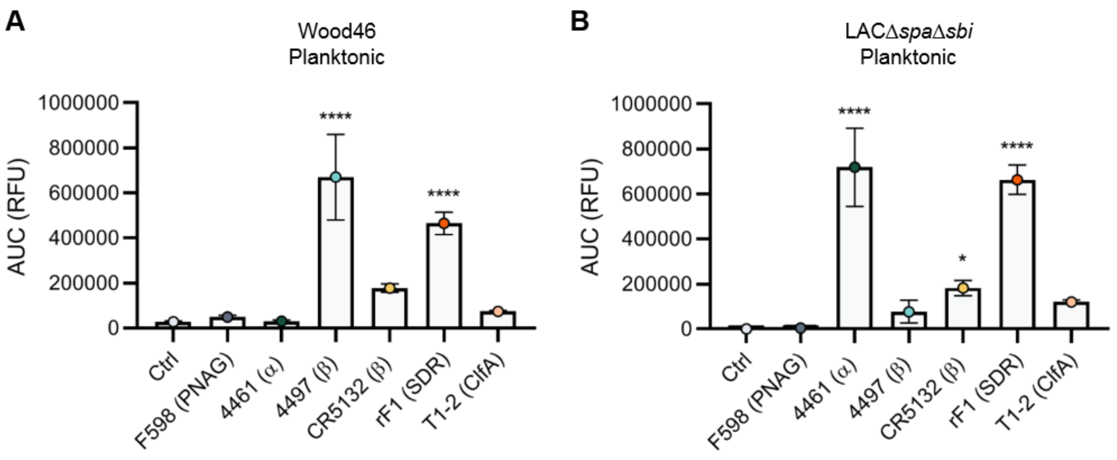


**Fig. S7.** Comparative binding of IgG1 mAbs to planktonic bacteria. Planktonic bacteria of Wood46 (A) and LACΔ*spa*Δ*sbi* (B) were grown to exponential phase and incubated with a concentration range of IgG1 mAbs. MAb binding was detected using APC-labeled anti-human IgG antibodies and flow cytometry. Data were expressed as AUC of the binding curve (mean + SD) relative to rF1-IgG1 of at least 3 independent experiments. One-way ANOVA followed by Dunnett test was performed to test for differences in antibody binding versus control and displayed only when significant as *P ≤ 0.05, **P ≤ 0.01, ***P ≤ 0.001, or ****P ≤ 0.0001.

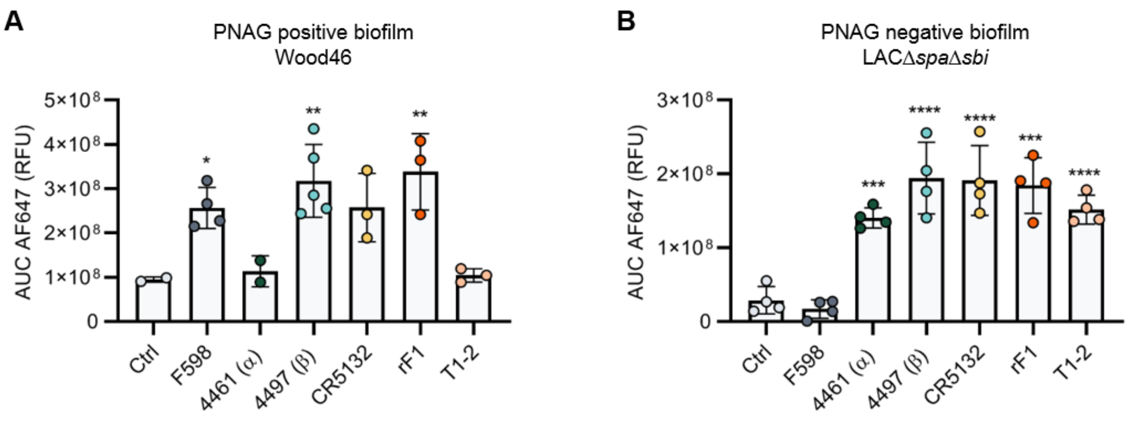


**Fig. S8.** Mean total fluorescence per Z-stack corresponds to plate reader data. The AF647 AUC of obtained Z-stack profiles of biofilms Wood46 (A) and LACΔ*spa*Δ*sbi* (B) was calculated with Prism 8.3.0. One-way ANOVA followed by Dunnett test was performed to test for differences in antibody binding versus control and displayed only when significant as *P ≤ 0.05, **P ≤ 0.01, ***P ≤ 0.001, or ****P ≤ 0.0001.

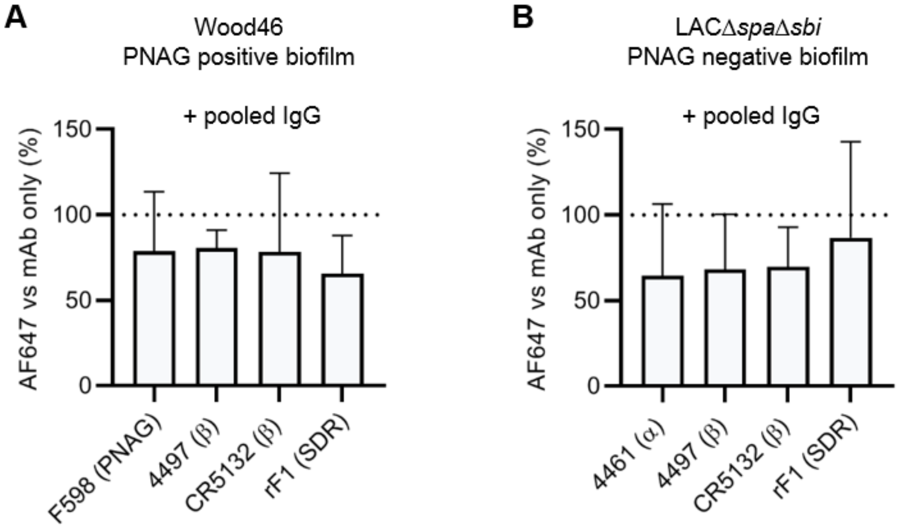


**Fig. S9.** Binding in presence of pooled serum IgG. Biofilm cultures of Wood46 (A) and LACΔ*spa*Δ*sbi* (B) were incubated with 10 µg/mL AF647-conjugated IgG1 mAbs in buffer or buffer containing 250 µg/mL pooled IgG. Data were expressed as AUC of the binding curve (mean + SEM) relative to rF1-IgG1 of at least 3 independent experiments. One-way ANOVA followed by Dunnett test was performed to test for differences in antibody binding versus control and displayed only when significant as *P ≤ 0.05, **P ≤ 0.01, ***P ≤ 0.001, or ****P ≤ 0.0001.

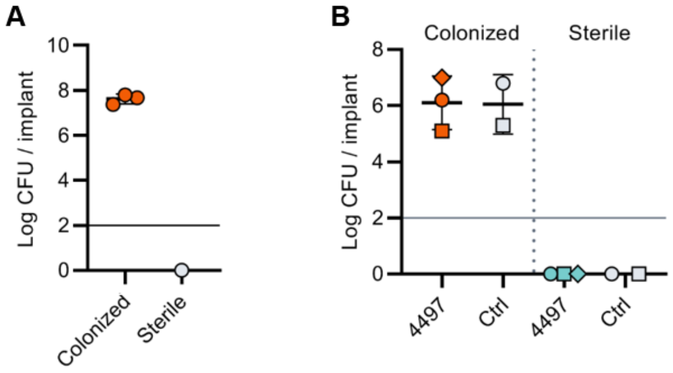


**Fig. S10.** CFU count before implantation and after implantation. (A) 5 mm PU catheter segments were inoculated with *S. aureus* LAC. After 48h of incubation, catheters were washed and sonicated and viable CFU counts recovered were determined. (B) Mice received subcutaneous pre-colonized and sterile catheters and 2 days later were injected with [$^{111}$In]In-4497-IgG1 or [$^{111}$In]In-Palivizumab. At time point 120 h, mice were sacrificed and catheters were removed to determine CFU counts. Horizontal lines indicate detection limit.

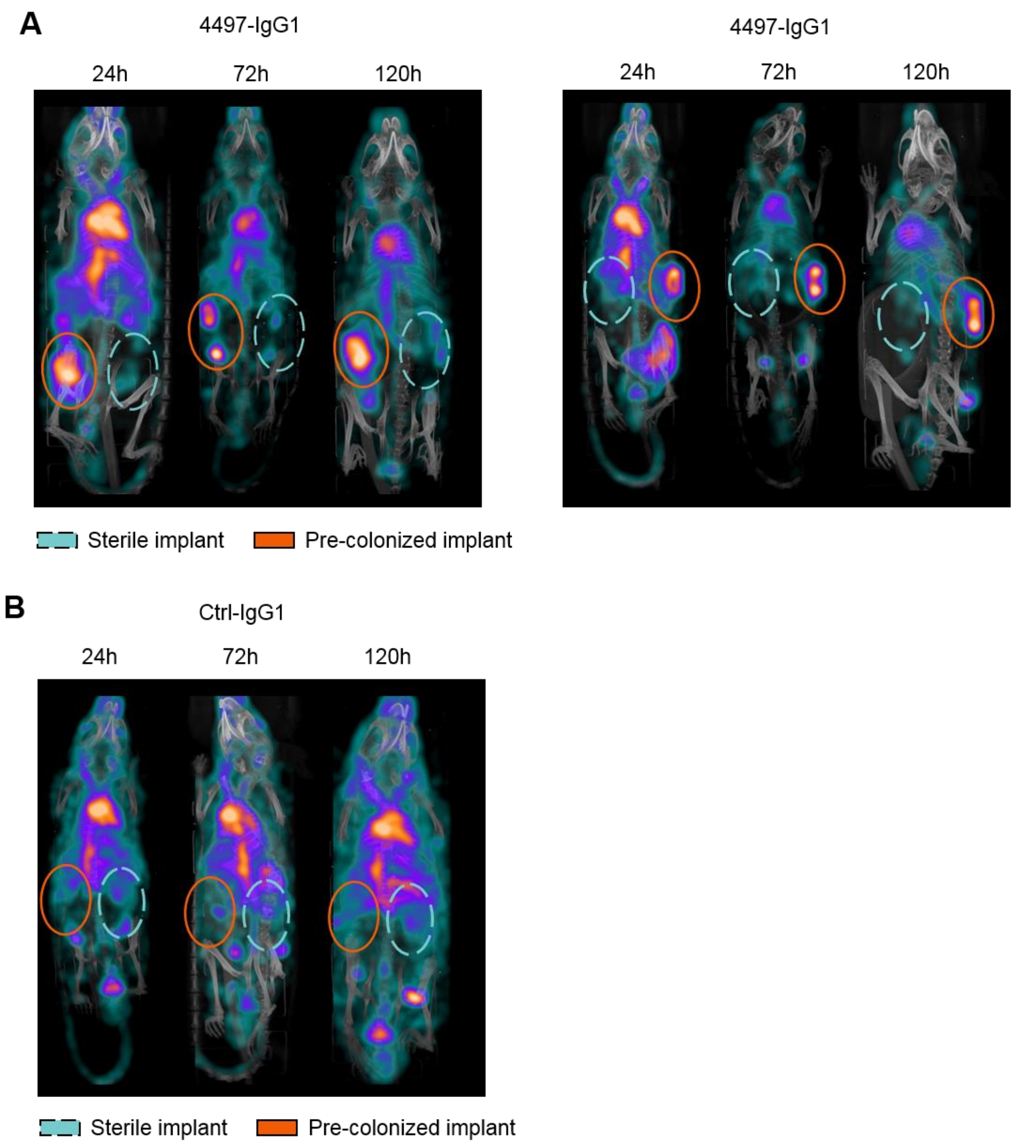


**Fig. S11.** Localization of [$^{111}$In]In-4497-IgG1 to subcutaneous implant infection in a mouse model. Maximum intensity projections of (A) [$^{111}$In]In-4497-IgG1 and (B) [$^{111}$In]In-Palivizumab injected in mice subcutaneously bearing pre-colonized (orange, full ellipses) and sterile (blue, striped ellipses) catheters.

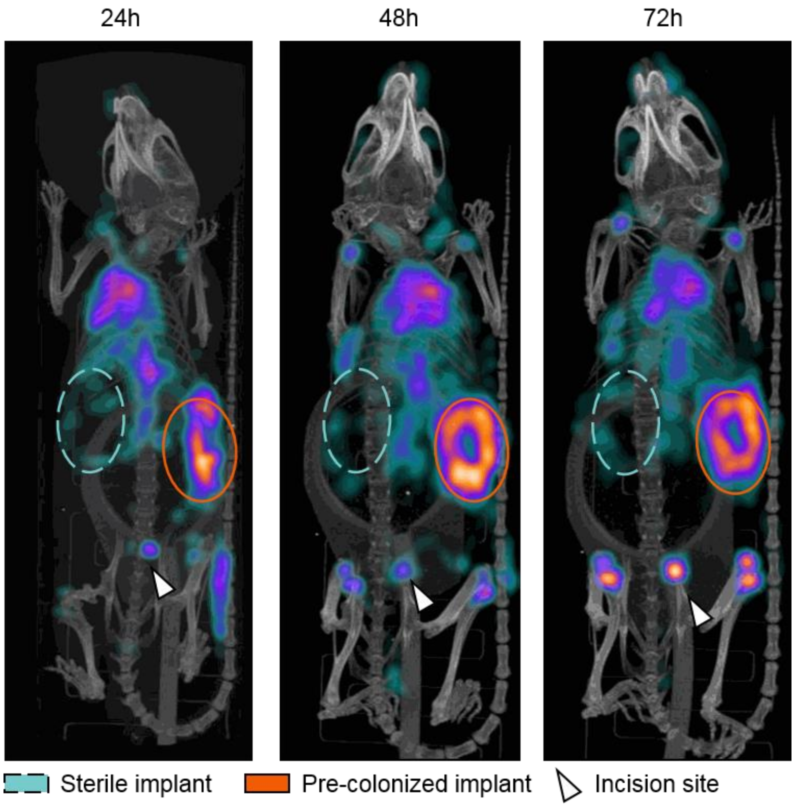


**Fig. S12.** Pilot study for localization of $[^{111}In]In$-4497-IgG1 to subcutaneous implant infection in a mouse model. One mouse received subcutaneous pre-colonized and sterile catheters and 2 days later was injected with 4MBq $[^{111}In]In$-4497-IgG1. The same mouse was imaged at 24 h, 48 h and 72 h. Maximum intensity projections of $[^{111}In]In$-4497-IgG1 injected in mice subcutaneously bearing pre-colonized (orange, full ellipses) and sterile (blue, striped ellipses) catheters.

**Table S1.** Protein sequences used for human monoclonal antibody production.

| **Clone, *target*** | **Sequence** | **Reference** |
|---|---|---|
| **VH variable heavy chain** | | |
| G2a2, *Anti-DNP* | DVRLQESGPGLVKPSQSLSLTCSVTGYSITNSYYWNWIRQFPGNKLEWMVYIGYDGSNNYNPSLKNRISITRDTSKNQFFLKLNSVTTEDTATYYCARATYYGNYRGFAYWGQGTLVTVSA | Gonzalez 2003 (3) |
| B12, *Anti-gp120* | QVQLVQSGAEVKKPGASVKVSCQASGYRFSNFVIHWVRQAPGQRFEWMGWINPYNGNKEFSAKFQDRVTFTADTSANTAYMELRSLRSADTAVYYCARVGPYSWDDSPQDNYYMDVWGKGTTVIVSS | Barbas 1993 (4)<br>Saphire 2001 (5) |
| 4461, *Anti-WTA(α)* | QVQLVQSGAEVRKPGASVKVSCKASGYSFTDYYMHWVRQAPGQGLEWMGWINPKSGGTNYAQRFQGRVTMTGDTSISAAYMDLASLTSDDTAVYYCVKDCGSGGLRDFWGQGTTVTVSS | WO/2014/193722 A1 |
| 4497, *Anti-WTA(β)* | EVQLVESGGGLVQPGGSLRLSCSASGFSFNSFWMHWVRQVPGKGLVWISFTNNEGTTTAYADSVRGRFIISRDNAKNTLYLEMNNLRGEDTAVYYCARGDGGLDDWGQGTLVTVSS. | WO/2014/193722 A1<br>Lehar 2015 (6)<br>Fong 2018 (7) |
| CR5132 | EVLESGGGLVQPGGSLRLSCSDSGFSFNNYWMTWVRQAPGKGLEWVANINRDGSDKYHVDSVEGRFTISRDNSKNSLYLQMNNLRADDAA VYFCARGGRTTSWYWRNWGQGTLVTVSS | US 2012/0141493 A1 |
| F598, *Anti-PNAG* | QVQLQESGPGLVKPSETLSLTCTVSGGSISGYYWSWIRQPPGKGLEWIGYIHYSRSTNSNPALKSRVTISSDTSKNQLSLRLSSVTAADTAVYYCARDTYYYDSGDYEDAFDIWGQGTMVTVSS | US/2006/0115486 A1 Seq25<br>Kelly-Quintos 2006 (8)<br>Soliman 2018 (9) |
| rF1, *Anti-GlcNac pan-SDR* | EVQLVESGGGLVQPGGSLRLSCAASGFTLSRFAMSWVRQAPGRGLEWVASINSGNNPYYARSVQYRFTVSRDVSQNTVSLQMNNLRAEDSATYFCAKDHPSSGWPTFDSWGPGTLVTVSS | WO/2016/090040 Seq13<br>Hazenbos 2013 (10) |
| T1-2, *Anti-ClfA* | QVQLKESGPGLVAPSQSLSITCAISGFSLSRYSVHWVRQPPGKGLEWLGMIWGGGNTDYNSALKSRLSISKDNSKSQVFLKMNSLQTDDTAMYYCARKGEFYYGYDGFVYWGQGTLVTVSA | WO 02072600 A2 |
| **VL variable light chain** | | |
| G2a2, *Anti-DNP* | DIRMTQTTSSLSASLGDRVTISCRASQDISNYLNWYQQKPDGTVKLLIYYTSRLHSGVPSRFSGSGSGTDYSLTISNLEQEDIATYFCQQGNTLPWTFGGGTKLEIK | Gonzalez 2003 (3) |

| | | |
|---|---|---|
| B12,<br>*Anti-gp120* | EIVLTQSPGTLSLSPGERATFSCRSSHSIRSRRVAWYQHKPGQ APRLVIHGVSNRASGISDRFSGSGSGTDFTLTITRVEPEDFALYY CQVYGASSYTFGQGTKLERK | Barbas 1993 (4)<br>Saphire 2001 (5) |
| 4461,<br>*Anti-WTA(α)* | DIQMTQSPDSLAVSLGERATINCKSSQSVLSRANNNYYVAWYQ HKPGQPPKLLIYWASTREFGVPDRFSGSGSGTDFTLTINSLQAE DVAVYYCQQYYTSRRTFGQGTKVEIK | WO/2014/193722 A1 |
| 4497,<br>*Anti-WTA(β)* | DIQLTQSPDSLAVSLGERATINCKSSQSIFRTSRNKNLLNWYQQ RPGQPPRLLIHWASTRKSGVPDRFSGSGFGTDFTLTITSLQAED VAIYYCQQYFSPPYTFGQGTKLEIK | WO/2014/193722 A1<br>Lehar 2015 (6)<br>Fong 2018 (7) |
| CR5132 | STDIQMTQSPSTLSASVGDRVTITCRASQSISSWLAWYQQKPG KAPKLLIYKASSLESGVPSRFSGSGSGTEFTLTISSLQPDDFATY YC QQYNSYPLTFGGGTKLEIK | US 2012/0141493 A1 |
| F598,<br>*Anti-PNAG* | QLVLTQSPSASASLGASVKLTCTLSSGHSNYAIAWHQQQPGKG PRYLMKVNRDGSHIRGDGIPDRFSGSTSGAERYLTISSLQSEDE ADYYCQTWGAGIRVFGGGTKLTVLG | US/2006/0115486 A1 Seq 26<br>Kelly-Quintos 2006 (8)<br>Soliman 2018 (9) |
| rF1,<br>*Anti-GlcNac pan-SDR* | DIQLTQSPSALPASVGDRVSITCRASENVGDWLAWYRQKPGKA PNLLIYKTSILESGVPSRFSGSGSGTEFTLTISSLQPDDFATYYC QHYMRFPYTFGQGTKVEIK | WO/2016/090040_ Seq14<br>Hazenbos 2013 (10) |
| T1-2,<br>*Anti-ClfA* | NIMMTQSPSSLAVSAGEKVTMSCKSSQSVLYSSNQKNYLAWY QQKPGQSPKLLIYWASTRESGVPDRFTGSGSGTDFTLTISSVQA EDLAVYYCHQYLSSYTFGGGTKLEIK | WO 02072600 A2 |
| **HC constant regions** | | |
| IgG1 | ASTKGPSVFPLAPSSKSTSGGTAALGCLVKDYFPEPVTVSWNS GALTSGVHTFPAVLQSSGLYSLSSVVTVPSSSLGTQTYICNVNH KPSNTKVDKKVEPKSCDKTHTCPPCPAPELLGGPSVFLFPPKPK DTLMISRTPEVTCVVVDVSHEDPEVKFNWYVDGVEVHNAKTKP REEQYNSTYRVVSVLTVLHQDWLNGKEYKCKVSNKALPAPIEK TISKAKGQPREPQVYTLPPSREEMTKNQVSLTCLVKGFYPSDIA VEWESNGQPENNYKTTPPVLDSDGSFFLYSKLTVDKSRWQQG NVFSCSVMHEALHNHYTQKSLSLSPGK | Kabat 1991 (11) |
| **LC constant regions** | | |
| Kappa class | RTVAAPSVFIFPPSDEQLKSGTASVVCLLNNFYPREAKVQWKVD NALQSGNSQESVTEQDSKDSTYSLSSTLTLSKADYEKHKVYAC EVTHQGLSSPVTKSFNRGEC | Kabat 1991 (11) |

**Movie S1 (separate file).** Localization of [$^{111}$In]In-4497-IgG1 to subcutaneous implant infection in a mouse model. 3D projections of [$^{111}$In]In-4497-IgG1 injected in mice subcutaneously bearing pre-colonized and sterile catheters. The same mouse was imaged at 24 h, 72 h and 120 h.

**Movie S2 (separate file).** 3D projections of [$^{111}$In]In-Palivizumab injected in mice subcutaneously bearing pre-colonized and sterile catheters. The same mouse was imaged at 24 h, 72 h and 120 h.
