## Supplementary figures and images for "Human monoclonal antibodies against *Staphylococcus aureus* surface antigens recognize *in vitro* biofilm and *in vivo* implant infections"

### Supplemental Movie 1

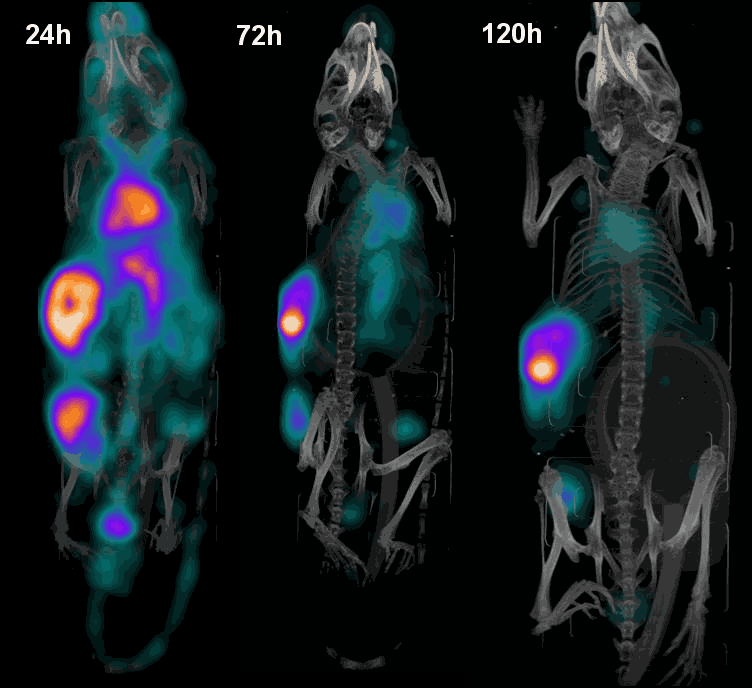

### Supplemental Movie 2

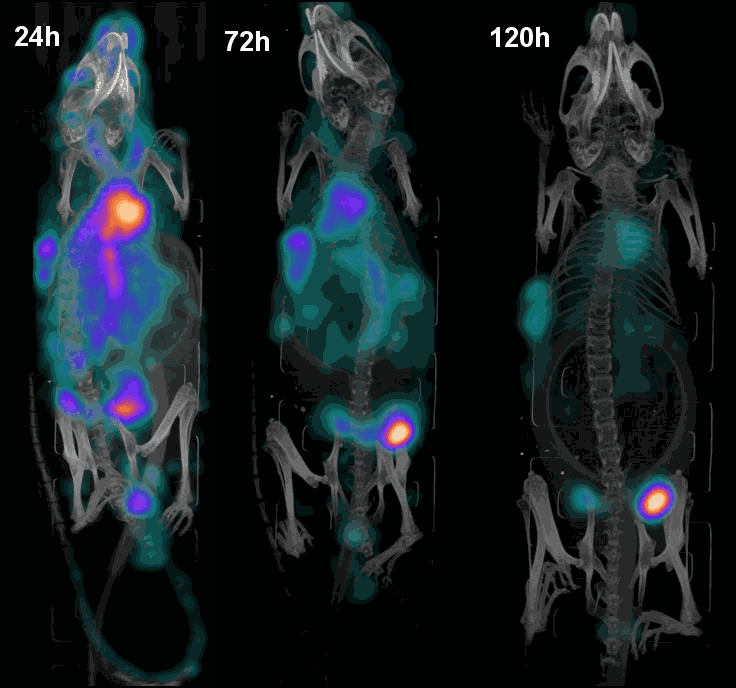
